# Pervasive transcription enhances the accessibility of H-NS-silenced promoters and generates bistability in *Salmonella* virulence gene expression

**DOI:** 10.1101/2022.04.26.489344

**Authors:** Nara Figueroa-Bossi, María Antonia Sanchez-Romero, Patricia Kerboriou, Delphine Naquin, Clara Mendes, Philippe Bouloc, Josep Casadesús, Lionello Bossi

## Abstract

In *Escherichia coli* and *Salmonella*, many genes silenced by the nucleoid structuring protein H-NS are activated upon inhibiting Rho-dependent transcription termination. This response is poorly understood and difficult to reconcile with the view that H-NS acts mainly by blocking transcription initiation. Here we have analysed the basis for the upregulation of H-NS-silenced *Salmonella* Pathogenicity Island 1 (SPI-1) in cells depleted of Rho-cofactor NusG. Evidence from genetic experiments, semi-quantitative 5’ RACE-Seq and ChiP-Seq shows that transcription originating from spurious antisense promoters, when not stopped by Rho, elongates into a H-NS-bound regulatory region of SPI-1, displacing H-NS and rendering the DNA accessible to the master regulator HilD. In turn, HilD’s ability to activate its own transcription triggers a positive feedback loop that results in transcriptional activation of the entire SPI-1. Significantly, single-cell analyses revealed that this mechanism is largely responsible for the coexistence of two subpopulations of cells that, although genetically identical, either express or don’t express SPI-1 genes. We propose that cell-to-cell differences produced by stochastic spurious transcription, combined with feedback loops that perpetuate the activated state, can generate bimodal gene expression patterns in bacterial populations.

## Introduction

Transcriptomic analyses of *E. coli* bacteria exposed to an inhibitor of transcription termination factor Rho have led to the recognition that a major activity of Rho and its cofactor NusG in growing cells is devoted to genome-wide suppression of ubiquitous antisense transcription genome-wide (1, 2). Thus, indirectly, these findings are also pertinent to transcription initiation, as they unveil the existence of a high-level spurious transcriptional noise apparently curbed by the termination activity of Rho. This notion gained momentum with the demonstration that *E. coli* genes contain a multitude of intragenic promoters in both sense and antisense orientations (3–5). The phenomenon is particularly dramatic in genomic regions thought to originate from horizontal transfer whose typically higher Adenine and Thymine (AT) content matches the sequence composition of the average bacterial promoter (6). The disproportionally high number of promoter-like sequences in AT-rich DNA can actually be a source of toxicity by causing RNA polymerase titration (7). The above studies have an additional common denominator: they both implicated the nucleoid structuring protein H-NS. On the one hand, the sites of intragenic Rho-dependent termination colocalized with the regions bound by H-NS (1); on the other hand, H-NS was shown to play the major role of silencing the spurious intragenic promoters (5). These findings renewed the interest in understanding the hidden complexity of H-NS-RNA polymerase interactions (8).

A small abundant protein, H-NS oligomerizes along the DNA upon binding to high affinity AT-rich nucleation sites and spreading cooperatively to adjacent sequences through lower affinity interactions (9–11). The oligomerization process generates a higher-order superhelical structure thought to contribute to DNA condensation (12). The resulting nucleoprotein filaments effectively coat the DNA and thereby hamper promoter recognition by RNA polymerase (13). In addition, H-NS can repress transcription through the formation of bridged or looped DNA structures that trap RNA polymerase in the open complex (14, 15) or act as roadblocks against transcript elongation (16, 17). In particular, bridged but not linear H-NS filaments have been shown to promote Rho-dependent transcription termination by increasing transcriptional pausing in vitro (17). H-NS has gained considerable attention since the discovery of its role as a xenogeneic silencer. Due to its affinity for AT-rich DNA, H-NS preferentially binds to, and prevents the expression of sequences acquired through horizontal transfer (18, 19). In doing so, H-NS protects the bacterium against the toxicity of foreign DNA (7, 20, 21) and allows the evolution of mechanisms for co-opting newly acquired functions and regulating their expression (22, 23). Indeed, the vast majority of H-NS-silenced genes are tightly regulated and expressed only under a limited set of conditions. Activation of H-NS-silenced genes typically results from the binding or the action of regulators able to displace H-NS (24–26). Unlike classical gene activation, H-NS counter-silencing exhibits considerable flexibility in the spatial arrangement of the regulator protein relative to the promoter (27).

In *Salmonella enterica*, H-NS silences most of the genes that contribute to virulence, including *Salmonella* Pathogenicity Islands (SPIs) that are specifically activated in the environment of the infected host (18, 19). SPI activation occurs in the form of a hierarchical and temporal regulatory cascade that begins with the expression of SPI-1, a 44 Kb-island encoding a Type III Secretion System (T3SS) that delivers effector proteins promoting intestinal colonization and epithelial cell invasion (28, 29). Several lines of evidence suggest that the process is initiated by HilD, a SPI-1-encoded AraC-type transcriptional regulator that activates the expression of a second master regulator, HilA, which in turns activates the T3SS along with the product of a third regulatory gene, *invF* (30). Acting as a hub integrating diverse environmental and physiological signals, HilD is itself regulated at multiple levels including mRNA stability (31) mRNA translation (32, 33) and protein activity (33, 34). However, central to the regulatory cascade is the ability of HilD to activate its own synthesis. HilD binds to an extended region upstream of the *hilD* promoter in vitro (35) and in vivo (36). The presence of this region among the DNA fragments bound by H-NS in chromatin immunoprecipitation (ChIP) experiments suggests that HilD stimulates transcription of its own gene by antagonizing H-NS (37). Interestingly, SPI-1 exhibits a bistable expression pattern characterized by the presence of two subpopulations of cells that either express or don’t express SPI-1 genes (38–43). In laboratory cultures, the SPI-1^OFF^ population vastly predominates; however, SPI-1^ON^ cells are continuously produced and persist for several generations (42, 44) despite the fitness cost associated with the synthesis and assembly of the T3SS, which results in growth retardation (45). Retarded growth, however, makes the SPI-1^ON^ subpopulation tolerant to antibiotics (46). SPI-1^OFF^ cells also benefit from bistability: inflammation triggered by the T3SS of SPI-1^ON^ cells leads to the production of reactive oxygen species in phagocytes. Such chemicals produce tetrathionate upon oxidation of endogenous sulphur compounds, and tetrathionate respiration confers a growth advantage to *Salmonella* over competing species of the intestinal microbiota (47). Furthermore, SPI-1^OFF^ cells can invade the intestinal epithelium, a capacity that may benefit the population as a whole by countering invasion by avirulent mutants (38, 39, 42).

We recently found that inhibiting Rho-dependent transcription termination, by mutation or through the depletion of Rho cofactor NusG, causes massive upregulation of many *Salmonella* virulence genes including all major SPIs (16). The magnitude and the span of these effects suggested that they were not produced locally but reflected the activation of a global regulatory response. This led us to turn our attention to HilD. The work described below confirmed the HilD involvement and provided new insight on the interplay between transcription elongation and bacterial chromatin. In particular, our data suggest that H-NS-bound regions are not completely impermeable to RNA polymerase. Occasional spurious transcription initiation events within these regions trigger a relay cascade whereby elongating transcription complexes, if not stopped by Rho, dislodge H-NS oligomers making more promoters accessible to RNA polymerase and regulatory proteins. In addition, these findings support a model for the mechanism underlying SPI-1 bistability.

## Results

### Most of the Salmonella response to NusG depletion is HilD-mediated

In all analyses described below, NusG depletion is achieved in strains with the sole copy of the *nusG* gene fused to a phage promoter under the control of an arabinose-inducible repressor (16). In the presence of arabinose (ARA), activation of the repressor gene causes the *nusG* gene to be turned off and its product to be progressively depleted. Although NusG is essential for *Salmonella* viability (48), the treatment is not lethal since residual NusG synthesis is sufficient to support growth. In fact, growth is nearly unaffected by ARA until bacteria enter early stationary phase. At this point the growth rate becomes significantly reduced, apparently as a side effect of the strong activation of SPIs (45).

To assess the possible role of HilD in the response to NusG depletion, we measured the expression of *lacZ* translational fusions to three SPI genes: *invB* (SPI-1) *sseE* (SPI-2) and *sopB* (SPI-5) in a *hilD^+^* strain and in a strain in which the *hilD* gene is replaced by a *tetRA* cassette. ARA exposure elicited a sharp increase in the expression of all three fusions in both *hilD*^+^ and *hilD*^-^ backgrounds; however, the changes in the *hilD^+^* strain occurred within a range between 10- to 50-fold higher than in the *ΔhilD::tetRA* mutant (Supplementary Fig. S1), pointing to the HilD involvement in the ARA-mediated activation of SPIs. To examine this response at the single cell level, we constructed in-frame fusions of superfolder green fluorescent protein (GFP^SF^) to HilD-regulated genes. Two fusions, in *hilA* and *invB*, were obtained by inserting the *gfp^SF^* open reading frame (orf) in the target gene; a third fusion, also in *hilA*, was made by concomitantly deleting a 28,266 bp segment spanning nearly the entire SPI-1 portion on the 3’ side of the fusion boundary. As initial experiments showed no significant differences in the behaviors the three strains, only the strain with the 28 Kb SPI-1 deletion (*hilA::gfp*^SF^ΔK28) was used for subsequent analyses.

### NusG depletion promotes SPI-1 bistability

Cells carrying *hilA*::*gfp*^SF^ΔK28 display a typical bistable phenotype characterised by the presence of two subpopulations of bacterial cells, of which only one subpopulation shows GFP fluorescence (Fig. *1A*). Significantly, growth in the presence of ARA causes the ratio between hilA^ON^ and hilA^OFF^ cells to increase dramatically in early stationary phase (Fig. 1*A*). Flow cytometric measurements show the increase to be more than 10-fold (Fig. 1*B*, Supplementary Fig. S2). SPI-1 bistability has been linked to the self-activating nature of *hilD* expression and it is thought to reflect cell-to-cell variability in HilD levels (32, 33, 41). In line with this model, *hilA*^ON^ cells are no longer detected in a strain carrying a 309 bp in-frame deletion removing the DNA binding motif in the carboxyl-terminal domain of HilD (Fig. 1*C*, Supplementary Fig. S2). These results suggest that NusG depletion allows a larger fraction of cells to reach the HilD autoactivation threshold. Consistent with this conclusion, RNA quantification by reverse transcription-quantitative polymerase chain reaction (RT-qPCR) shows that the ARA treatment causes a large increase in *hilD* transcription when HilD is functional but a smaller increase in the *hilDΔ309* mutant (Fig. 2*A*). The analysis reveals that HilD is also needed for the expression of the adjacent *prgH* gene (Fig. 2*B*). At first sight, this may seem surprising since *prgH* is thought to be activated by HilA, not by HilD (37, 49) and the strain used in Fig. 2*B* carries the *hilA::gfp*^SF^ΔK28 allele which removes over two thirds of the *hilA* sequence. However, we note that the fusion retains the N-terminal 112 amino acids (aa) domain of HilA previously implicated in *prgH* promoter recognition (50) suggesting that the HilA-GFP chimera retains the ability to activate *prgH*.

**Figure 1.**
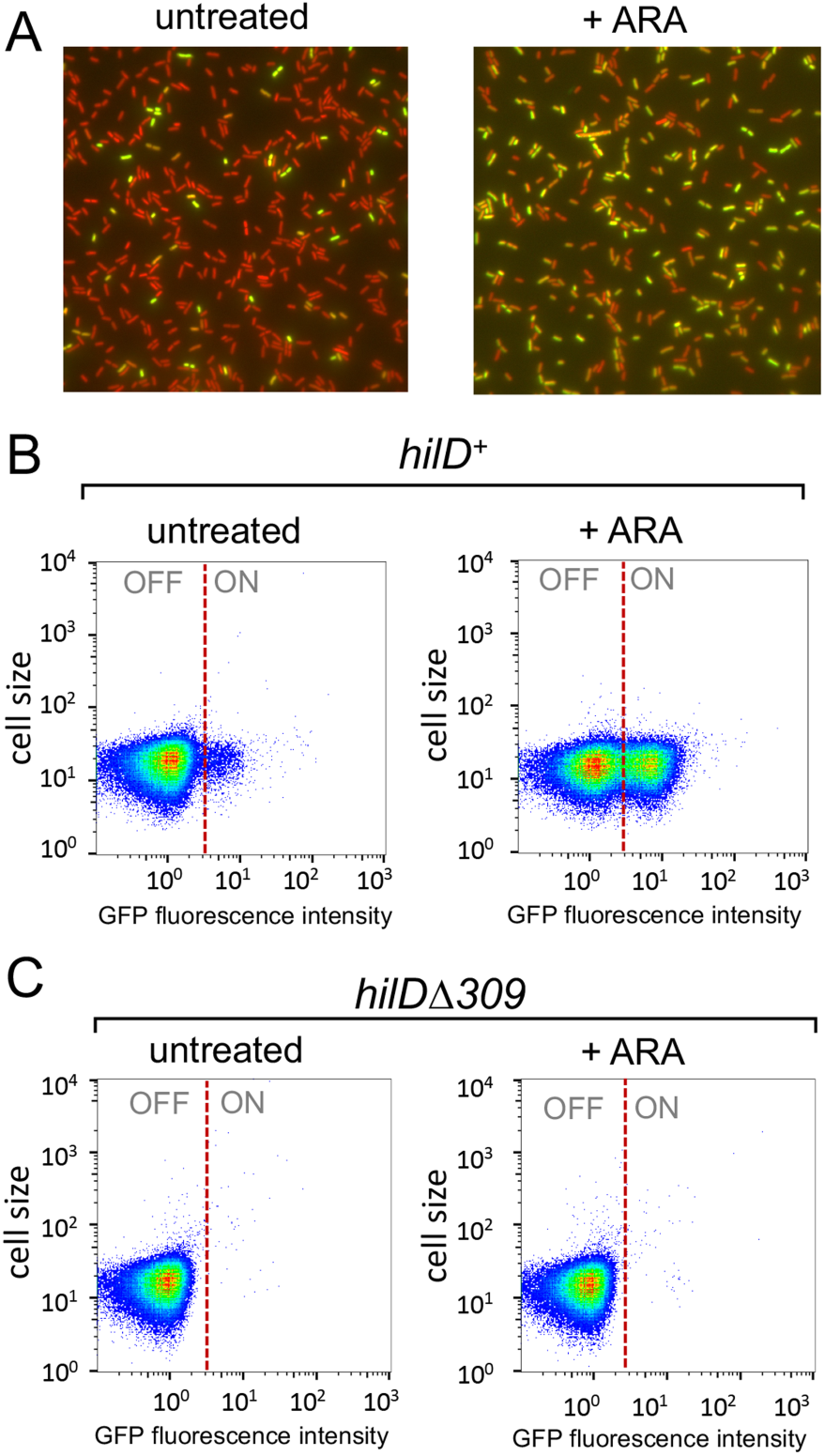
NusG depletion enhances HilD-dependent SPI-1 bistability. The strains used, MA14302 (*hilD^+^*) and MA14561 (*hilDΔ309*), carry a *hilA-gfp*^SF^ translational gene fusion (*hilA*::*gfp*^SF^ΔK28) and a chromosomal P^Tac^ promoter-mCherry gene fusion in the ARA-inducible-NusG depletion background. (*A*) Representative image of MA14302 cells grown at 37°C to early stationary phase visualized by fluorescence microscopy under 100x magnification. (*B*) and (*C*) Representative flow cytometry analysis of cells from strains MA14302 (*B*) and MA14561 (*C*) grown as in (*A*). The GFP fluorescence intensity distribution was examined in strains carrying *gfp* translational fusions. The full genotypes of MA14302 and MA14561 are shown in *Supplementary* Table S1. For the construction of *hilA::gfp*^SF^ΔK28 and *hilD*Δ*309* by λ *red* recombineering, DNA primers (listed in *Supplementary* Table S2) were used as detailed in *Supplementary* Table S3.

**Figure 2.**
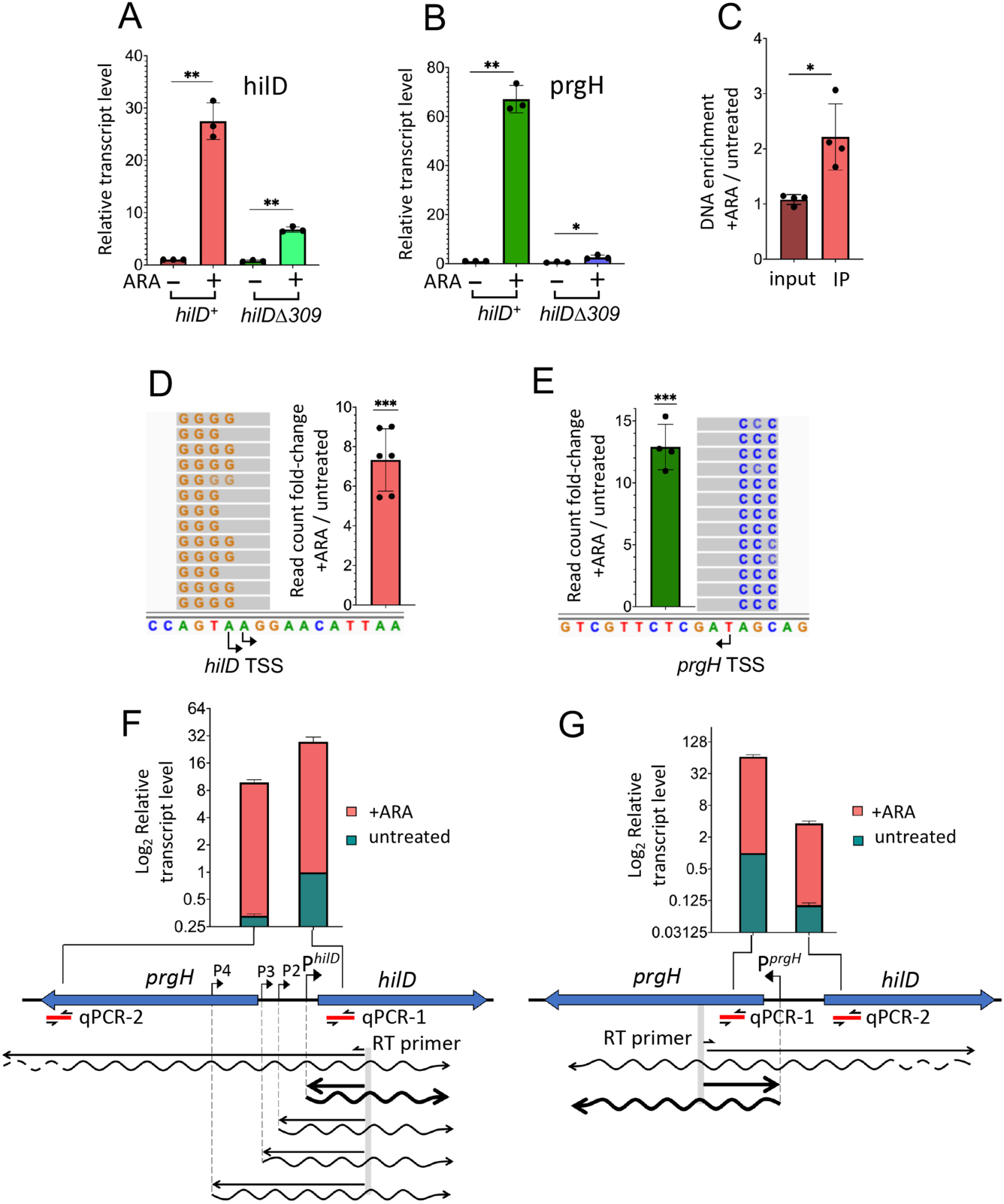
NusG depletion induces HilD-dependent activation of *hilD* and *prgH* promoters. (*A*) and (*B*) Quantification of *hilD* mRNA (*A*) and *prgH* mRNA (*B*) from strains MA14302 (*hilD*^+^) and MA14561 (*hilD*Δ*309*) grown to early stationary phase in the absence or in the presence of 0.1% ARA. RNA was quantified by two-step reverse transcription-quantitative PCR (RT-qPCR). Ct values were normalized to the Ct values determined for *ompA* mRNA. Transcript levels are shown relative to those of untreated MA14302, set as 1. (*C*) Measurement of HilD protein binding to the *hilD* promoter. HilD-bound DNA was isolated by chromatin immunoprecipitation from strain MA14363 (carrying a chromosomal *hilD*-3xFLAG fusion) and quantified by real-time PCR (ChIP-qPCR). Ct values were normalized to the values of a *katE* gene reference. Results are presented as ratios between the values measured in cells grown in ARA-supplemented medium and the values from untreated cells. (*D*) and (*E*) 5’ RACE-Seq analysis of *hilD* and *prgH* promoter activity, respectively. RNA from strain MA14302 was reverse-transcribed in the presence of a template-switching oligonucleotide (TSO). The resulting cDNA was used as template for semiquantitative PCR with primers carrying Illumina adapter sequences at their 5’ ends. Amplified DNA was subjected to high-throughput sequencing. Read counts were normalized to those measured at the *ompA* promoter. Results shown represent the ratios between the normalized counts from ARA-treated cells and those from untreated cells. (*F*) and (*G*) Contribution of distal transcription to *hilD* and *prgH* RNA levels, respectively. RNA was reverse-transcribed with primers annealing inside the promoter-proximal portion of *hilD* or *prgH* (predicted transcripts are depicted as wavy lines). The resulting cDNAs (straight lines) were used for qPCR amplification with primers annealing close to (qPCR-1) or farther away from (qPCR-2) the RT priming site. Ct values were normalized to the Ct values determined for *ompA* mRNA. Transcript levels are shown relative to those of untreated MA14302 cells, set as 1. All the data in this figure originate from ≥3 independent experiments (with error bars indicating standard deviations). Statistical significance was determined by unpaired two-tailed Student t-tests with Welch’s correction for unequal variances (*, P ≤ 0.05; **, P, ≤ 0.01; ***, P ≤ 0.001). In (*F*) and (*G*), the calculated P values for the differences between untreated samples (green bars) were 0.0002 (*F*) and < 0.0001 (*G*). The P values for the ARA-treated samples (red bars) were 0.0108 (*F*) and 0.0025 (*G*). The oligonucleotides used as primers in the above experiments are listed in Supplementary Table S4. Further experimental details are provided in Materials and Methods.

### Pervasive transcription activates the hilD promoter

The central role of HilD in the response to NusG depletion was further corroborated by the observation that ARA treatment stimulates HilD binding to the *hilD* promoter region (Fig. 2*C*), a response that correlates with the activation of the *hilD* and *prgH* promoters (Fig. 2*D* and *E*). This last set of data were obtained performing semi-quantitative 5’ rapid amplification of cDNA ends (5’ RACE) generated by template switching reverse transcription (TS RT) (51). In this method, reverse transcription is primed by a gene-specific primer and carried out in the presence of a template switching oligonucleotide (TSO). The 5’ ends of RNAs are defined by the position the cytidine repeats (typically 3 or 4) that are added by reverse transcriptase when it reaches the end of the RNA and switches to the TSO (51). Use of primers carrying Illumina adaptors for the PCR step allows for the analysis to be performed by high throughput sequencing (RACE-Seq). Here, primers were designed to detect transcription initiation taking place at the primary *hilD* and *prgH* promoters as well as at three secondary promoters previously identified by Kröger and coworkers (52) (Supplementary Fig. S3). Read summarization at each of the transcription start sites (TSSs) showed that ARA exposure causes the number of transcripts initiating at *hilD* and *prgH* primary TSSs to increase 7-fold and 13-fold, respectively (Fig. 2*D* and *E*). The most likely explanation of this effect is that NusG depletion allows transcription complexes formed outside the *prgH-hilD* promoter region to invade this region displacing H-NS and triggering HilD autogenous activation. Prime suspects for this effect are the secondary *hilD* promoters whose activity contributes to the *hilD* mRNA increase (Supplementary Fig. S4*A*). However, the experiment does not distinguish whether the increase in the number of reads associated with the secondary TSSs is due to a larger proportion of transcripts reaching the RT primer site (and thus susceptible to reverse transcription) in ARA treated cells or reflect an increase in promoter activity. If the latter were true, the implication would be that the secondary promoters are themselves activated by transcription initiating elsewhere, presumably further upstream, and they relay the effect to the *hilD* promoter. To address this point, we performed parallel qPCR measurements using primers pairs annealing at proximal and distal positions relative to the RT priming site. Results showed that a significant fraction of transcripts entering the *hilD* coding sequence in ARA-treated cells initiate as far as over 1400 bp upstream from the *hilD* promoter (compare red bars between PCR-1 and PCR-2 in Fig. 2*F*), thus considerably upstream relative to the secondary promoters. It is therefore conceivable that the elongation of these overlapping transcripts may activate the secondary promoters. Interestingly, this class of transcripts is also detectable in untreated cells (green bars in Fig. 2*F*) albeit at very low level. A similar trend is observed in the opposite strand where a fraction of *prgH* RNAs originates from antisense *hilD* transcription (Fig. 2*G*).

To further assess the contribution of upstream transcription to *hilD* promoter activity, the SPI-1 segment extending from the left boundary of the island up to a position 610 bp upstream of the *hilD* main TSS was deleted and replaced by a cassette comprising the *tetR* gene and the TetR-repressed P^tetA^ promoter (Fig. 3*A*). The transcriptional responses to ARA and to the P^tetA^ inducer, Anhydrotetracycline (AHTc), alone or combined, were analyzed by RT-qPCR and 5’RACE Seq. ARA was still able to activate transcription of *hilD* and *prgH* in the new background (Fig. 3*B* and *C*); however, activation was more moderate than seen above with the parental strain (compare to Fig. 2*A* and *B*, respectively); furthermore, no significant changes were observed at the level of the *hilD* and *prgH* primary TSSs (Fig. 3*D* and *E*). These findings corroborate the idea that the region deleted in the new construct contributes

**Figure 3.**
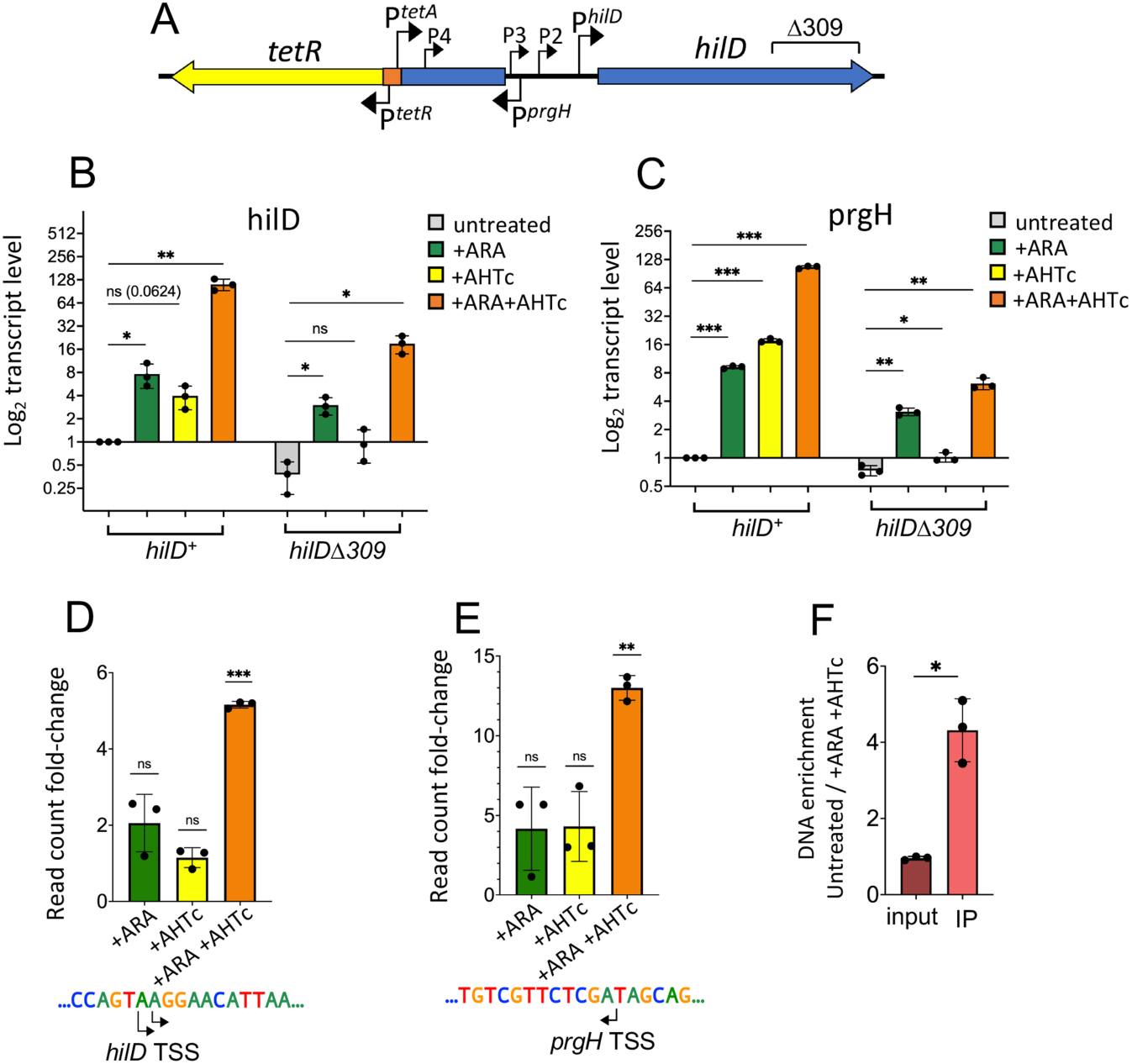
Overlapping transcription triggers HilD-dependent activation of *hilD* and *prgH* promoters in NusG-depleted cells. (A) Schematic diagram showing the gene organization on the 5’ side of *hilD* in strains MA14358 (*hilD*^+^) and MA14569 (*hilDΔ309*). Both strains contain a *tetR-P*^tetA^ cassette that replaces a 10,828 bp segment of SPI-1 and places the P^tetA^ promoter 610 bp upstream of the main *hilD* TSS. (B) and (C) Quantification of *hilD* mRNA (B) and *prgH* mRNA (C) in cells grown to early stationary phase in the presence or absence of either ARA or AHTc, or in the presence of both. RNA was quantified by two-step RT-qPCR. Ct values were normalized to the Ct values determined for *ompA* mRNA. Transcript levels are shown relative to those of untreated MA14358, set as 1. (*D*) and (*E*) 5’ RACE-Seq analysis of *hilD* and *prgH* promoter activity, respectively. RNA from strain MA14358 grown under the different conditions was processed as described in the legend of Fig. 2*D* and *E*. Results shown represent the ratios between the normalized read counts from treated cells and those from untreated cells. (*F*) Measurement of HilD protein binding to the *hilD* promoter. HilD-bound DNA was isolated by chromatin immunoprecipitation from strain MA14505 (carrying a chromosomal *hilD*-3xFLAG fusion) and quantified by real-time PCR (ChIP-qPCR). Ct values were normalized to the values of a *katE* gene reference. Results are presented as ratios between the values measured in cells grown in a medium supplemented with ARA and AHTc and those from untreated cells. All the data in this figure originate from three or more independent experiments (with error bars indicating standard deviations). Statistical significance was determined by unpaired two-tailed Student’s t-tests with Welch’s correction for unequal variances (ns, P> 0.05; *, P ≤ 0.05; **, P ≤ 0.01; ***, P ≤ 0.001). The oligonucleotides used as primers in the above experiments are listed in Supplementary Table S4.

to the amplitude of the ARA effects. Activating P^tetA^ with AHTc stimulates *hilD* and *prgH* transcription (Fig. 3*B* and *C*); here too, however, the effect remains limited and undetectable by 5’RACE-Seq at the primary TSSs (Fig. 3*D* and *E*). In contrast, when AHTc and ARA are used conjointly, transcription of both *hilD* and *prgH* genes is strongly activated (Fig. 3*B* and *C*) and an initiation burst is observed at the level of both primary promoters (Fig. 3*D* and *E*). Significantly, this burst correlates with increased occupancy of the *hilD* promoter by the HilD protein (Fig. 3*F*). The secondary promoters exhibit a similar overall response, which is however characterized by pronounced scatter in the individual Ara/AHTc treatments (Supplementary Fig. S4*B*). The overall strength of the *hilD* response can be correlated with the detection of higher levels of P^tetA^ transcription in the presence of ARA (Supplementary Fig. S4*C*).

In the background of the *hilA-gfp^SF^* fusion (which deletes the right two thirds of SPI-1), the deletion generated by the *tetR*-P^tetA^ insertion constitutes a minimal system with only two SPI-1 genes, *hilD* and *pphB*, remaining intact. Significantly, this strain still exhibits the HilD-dependent bistable phenotype, suggesting that HilD is not only required but also sufficient for bistability (Fig. 4*A* and *B*). Growth in a medium supplemented with AHTc affects the basal *hilA*^ON^/*hilA*^OFF^ ratio only marginally unless ARA is also present, in which case the vast majority of the cell population switches to the *hilA^ON^* status (Fig. 4*A* and *B*, Supplementary Fig. S5).

**Figure 4.**
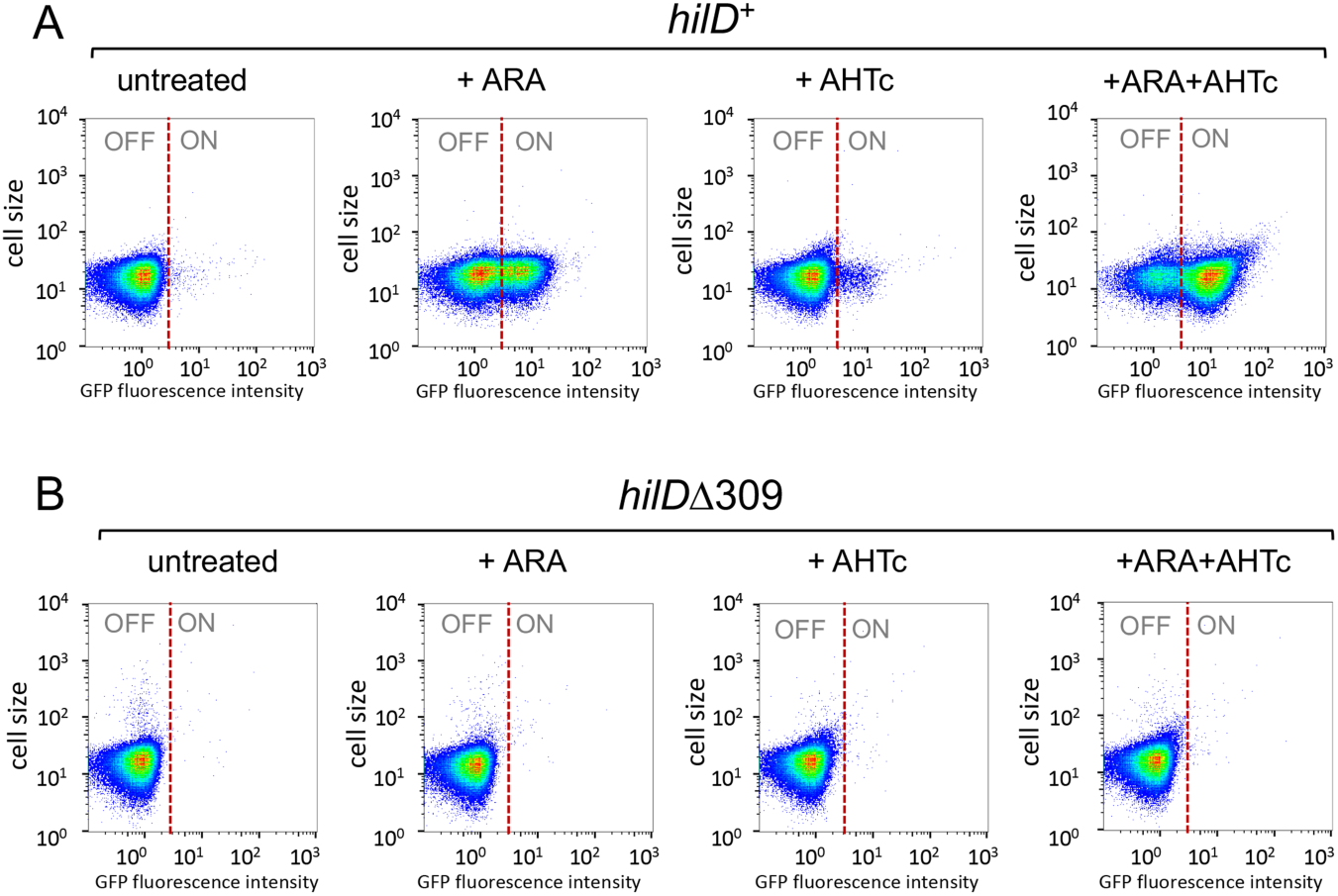
Overlapping transcription promotes HilD-dependent bistability of SPI-1 expression. Strains MA14358 (*hilD*^+^) and MA14569 (*hilD*Δ*309*), carry the *tetR*-P^tetA^ cassette (Fig. 3*A*) combined with *hilA*::*gp*^SF^ΔK28 and the chromosomal P^Tac^-*mCherry* in the ARA-inducible-NusG depletion background. Cells grown to early stationary phase under the indicated conditions were used for single-cell analysis by flow cytometry. GFP fluorescence was measured and the distribution of *hilA*^OFF^ and *hilA*^ON^ cells is shown in heat maps. (*A*) MA14358. (*B*) MA14569.

To confirm that the ARA effects depend on transcription originating upstream of *hilD*, and not by an alternative, unidentified mechanism activating *hilD* expression when NusG is depleted, we constructed a strain carrying the strong Rho-independent transcription terminator from the histidine operon attenuator region (Term^hisL^) immediately downstream from P^tetA^ in the *hilA::gfp*^SF^ΔK28 background (Supplementary Fig. S6*A* and *B*). During construction, a clone displaying strong green fluorescence on a plate supplemented with ARA and AHTc was identified. Sequence analysis revealed that this isolate harbors a deletion removing 6 out of the 9 repeated Us at the 3’ end of Term^hisL^. Both the strain with the wild-type Term^hisL^ insert and the Δ6U derivative were used for bistability assays. Results showed that Term^hisL^ abolishes all effects of ARA on *hilA* OFF/ON ratios both in the absence and in the presence of AHTc (Supplementary Fig. S6*C* and *E*). Interestingly, the Δ6U deletion reverses this pattern causing about half of the cell population to switch to the *hilA* ON status in the presence of ARA alone and virtually the entire population to switch ON in the presence of ARA and AHTc combined. (Supplementary Fig. *S6D* and *F*). These results provide conclusive evidence that transcription originating more than 600 bp upstream of *hilD’s* primary TSS is solely responsible for the effects of NusG depletion on *hilD* expression. Since *hilD* secondary promoters are all located downstream from Term^hisL^, these data also support the idea that they play no direct role in the ARA-induced activation of the primary promoter.

### Viewing SPI-1 upregulation at the chromatin level

In parallel with the above studies, we sought to determine whether NusG depletion affected the binding of H-NS to SPI-1 and other genomic islands. For this purpose, chromatin immunoprecipitation coupled with high-throughput sequencing (ChIP-Seq) was performed in strains carrying the NusG-repressible allele and an epitope tagged version of H-NS. Examination of the ChIP-Seq profiles in the SPI-1 section of the genome showed a succession of peaks and valleys consistent with the presence of multiple contiguous patches of oligomerized H-NS separated by segments with little or no H-NS bound (Fig. 5*A*). Superimposing the profiles from cells growing in the absence or in the presence of ARA reveals small but nonetheless appreciable differences in the levels of DNA fragments bound by H-NS. One can see that a number of peaks shrink as a result of the ARA treatment (Fig. 5*A*). Interestingly, the most conspicuous changes are detected in the *hilD-hilA* ad *invF-invH* sections of SPI-1, corresponding to the locations of main regulatory hubs (30). In contrast, no changes are observed at the far-right end of SPI-1 (*pigA-pphB* segment (53)) and in the central portion of the island.

**Figure 5.**
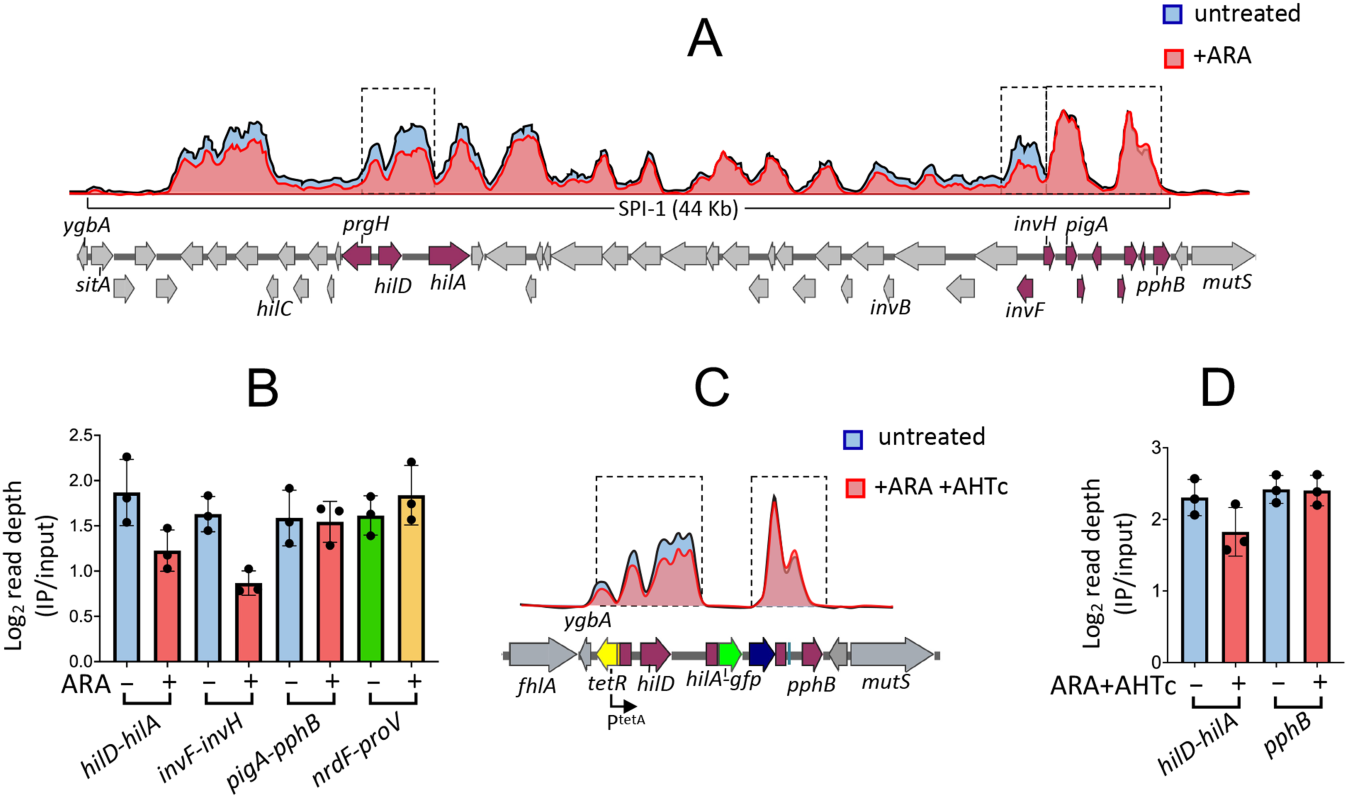
NusG depletion affects H-NS binding to specific portions of SPI-1. (*A*) Representative ChIP-Seq profiles from NusG-depletable strain MA13748 (*hns*-3xFLAG) grown to early stationary phase in the presence or absence of 0.1% ARA. (*B*) Read depth quantification in the sections framed by the dashed rectangles in (*A*) and in the *nrdF-proV* intergenic region. Read depth values (determined by the bedcov tool of the Samtools suite) were normalized to the values from the entire genome. The results shown represent the ratios between the normalized values from IP samples and those from input DNA. (*C*) Representative ChIP-Seq profiles from NusG-depletable strain MA14513 (Δ[*sitA-prgH*]::*tetR*-P^tetA^, *hilA*::*gfp*^SF^ΔK28, *hns*-3xFLAG) grown to early stationary phase in the presence or absence of ARA + AHTc. (*D*) Read depth quantification in the intervals framed by the dashed rectangles in (*C*). Read depth was calculated and normalized as in (*B*). The data in (*B*) and (*C*) represent the means from three independent ChIP-Seq experiments (with error bars indicating standard deviations).

Read depth quantification confirmed the profile changes. ARA treatment lowers H-NS binding in the *hilD-hilA* and *invF-invH* intervals by 34% and 47%, respectively, while having no effect in the *pigA-pphB* region (Fig. 5*B*). Likewise, no appreciable differences are observed at the *proV* locus (54) (Fig. 5*B*). ChIP-Seq analysis was also performed in the strain carrying the *tetR*-P^tetA^- and *hilA*::*gfp*^SF^ΔK28-associated deletions, comparing unchallenged cells to cells grown in the presence of both ARA and AHTc (Fig. 5*C*). Somewhat surprisingly, the double treatment caused only a 20% reduction of H-NS binding in the *hilD-hilA* interval (Fig. 5*D*). In both above analyses, visual inspection of the profiles around other H-NS-bound loci known to be upregulated in NusG-depleted cells (16) failed to reveal appreciable differences. Finding that transcriptional changes produce comparatively small or undetectable alterations in H-NS binding is not novel (24, 26) and suggests that H-NS-DNA complexes exist dynamically and rapidly reform after the passage of transcription elongation complexes.

## Discussion

This study was aimed at understanding why impairing Rho-dependent transcription termination by depletion of Rho-cofactor NusG relieves H-NS silencing of *Salmonella* pathogenicity islands. We show that NusG depletion triggers a positive feedback loop that generates and maintains HilD, the master regulator of the *Salmonella* virulence regulatory cascade. Accumulation of HilD is primarily responsible for H-NS counter-silencing in NusG-depleted cells. This is directly demonstrated for SPI-1, SPI-2 and SPI-5 genes, but it seems likely that the HilD involvement may extend to most, if not all, islands and islets found upregulated in NusG-depleted cells in a previous study (16). Note that although SPI-1 and SPI-2 are generally activated in response to sharply different cues, a HilD-mediated crosstalk allows expression of SPI-2 genes under conditions unusual for this island, notably in rich medium (55, 56).

By oligomerizing along the DNA, H-NS silences not only *bona fide* promoters at the 5’ end of genes but also a plethora of spurious intragenic promoters that “infest” A/T-rich horizontally acquired DNA (5–7). Finding that the inhibition of Rho or NusG causes widespread sense and antisense transcription of H-NS-silenced genes (1, 16) suggests that H-NS-bound DNA is susceptible to transcriptional invasion and that Rho (recruited by NusG) acts to prevent elongation of invading transcription complexes. Various lines of evidence suggest that H-NS-bound regions are not totally impermeable to RNA polymerase. Existence of several very short transcripts initiating from within H-NS-associated loci was previously inferred from a genome-wide analysis of TSSs in *E. coli* (4). More recently, parallel ChIP-Seq TSS mapping experiments showed a clear TSS being used upstream of the *E. coli ydbCD* operon, even when H-NS was present, but no full-length mRNA (6). Finally, in our previous work, we found that a *tetR*-P^tetA^ cassette placed only 57 bp away from the H-NS nucleation site in the *leuO* promoter region of *Salmonella* normally responds to AHTc induction (although leading to LeuO synthesis only when NusG is depleted) (16). Elongating through a patch of oligomerized H-NS, RNA polymerase can dislodge H-NS and allow other RNA polymerase molecules to gain access to normally silenced promoters thus further contributing to transcriptional noise (57, 58). The data presented here show that transcriptional “noise” can be converted into a true regulatory “melody” if the activated promoter directs the synthesis of a positive autoregulator. In the model schematized in Fig. 6, we posit that a transcription complex formed at a spurious promoter (“Px”), if not stopped by Rho, “unzips” the H-NS nucleoprotein filament in the *hilD* promoter region, triggering a positive feedback loop that results in HilD accumulation and concomitant derepression of both *hilD* and *prgH*. Antisense transcription from inside *hilD* (not shown for simplicity in Fig. 6) may contribute to destabilization of the H-NS-DNA complex. Note that transcription may not need to travel all the way to the promoter sequence in order to cause H-NS dissociation. Due to the multi-contact nature of the H-NS-DNA interaction (54), disruption of contacts at the edge of the oligomerization patch could be sufficient to destabilize the entire patch. By linking *hilD* activation to a stochastic and likely infrequent transcription event – *i.e*., initiation at a spurious promoter or readthrough of a Rho-dependent terminator by a spurious transcript – the model can explain the bistability in the expression of HilD-regulated loci and suggests that the frequency of these events may set SPI-1 ON/OFF subpopulation ratios. The HilD/H-NS interplay in the regulation of SPI-1 bears analogy with the mechanism regulating the expression of the Locus of Enterocytes Effacement (LEE) of enteropathogenic *Escherichia coli*. Here too, expression is characterized by a bistable response (59), suggesting that the interplay between H-NS and regulatory proteins (Ler in this case) may constitute an elemental premise for bistability. Whether LEE regulation responds to pervasive transcription is currently unknown.

**Figure 6.**
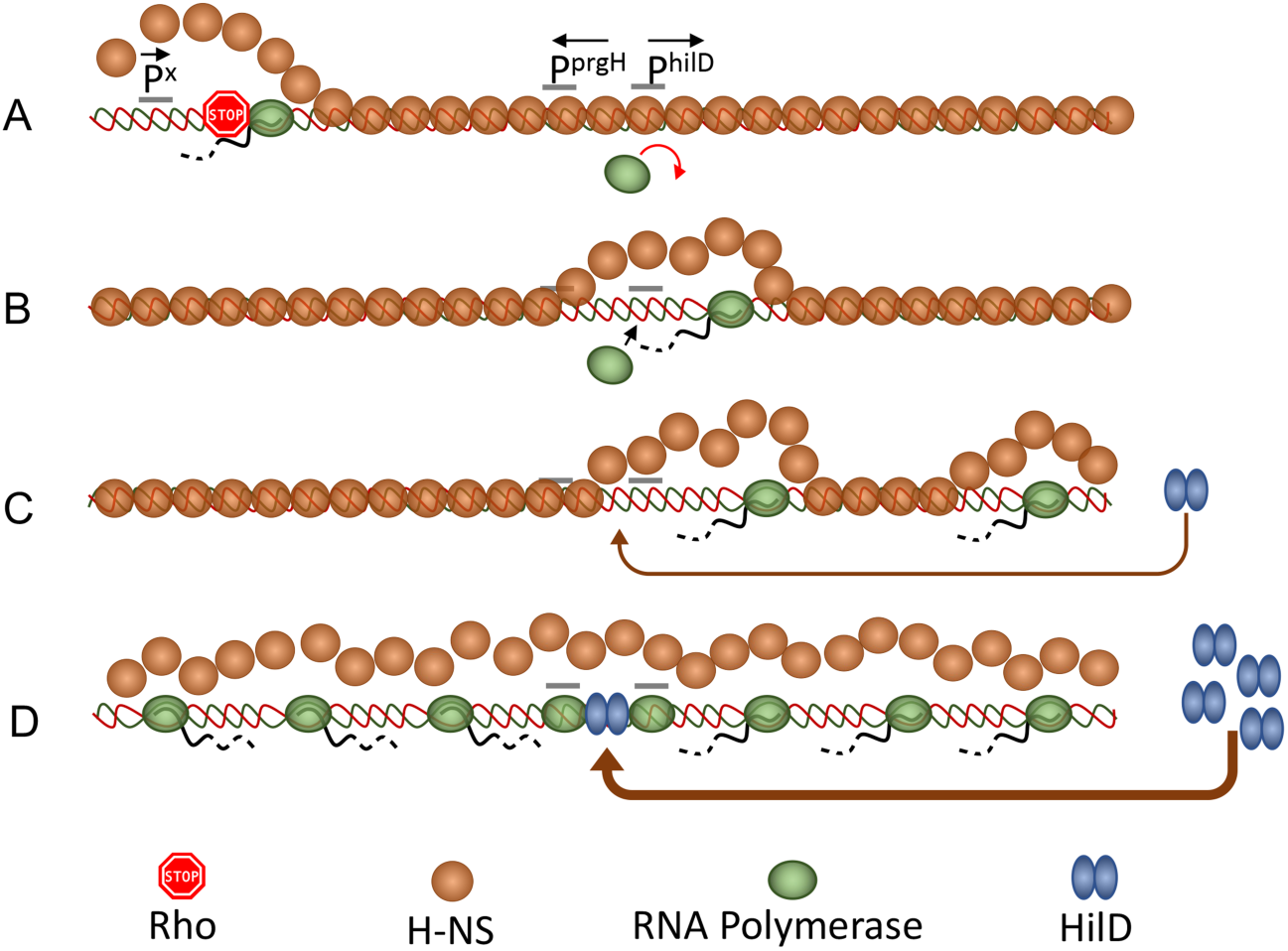
Model for activation of *hilD* and *prgH* promoters by overlapping transcription. (*A*) A spurious transcription initiation event occurs at the edge of a patch of oligomerized H-NS (orange circles). Transcript elongation through bound H-NS is prevented by NusG-mediated recruitment of Rho factor (stop sign). (*B*) Occasionally, the transcript eludes Rho termination and progresses along the DNA dislodging H-NS in front of its path. This action opens a kinetic window during which RNA polymerase (green ovals) can bind to promoters that become exposed, including *hilD* secondary promoters (not shown) and the primary *hilD* promoter. (*C*) Activation of the *hilD* promoter leads to an increase in the levels of HilD protein (blue double-ovals), which, upon binding to the *hilD* regulatory region, further stimulates *hilD* transcription and protein production. (*D*) This locks the system in a positive feedback loop: accumulation of HilD leads to more *hilD* transcription and more HilD protein made. Through HilA (not shown) it also results in high-level transcription of the *prgH* gene. Divergent transcription further enhances the accessibility of additional spurious promoter sequences further contributing to runaway transcription activation.

Eukaryotic genomes, including the predominant non-coding fraction of human genomes, are pervasively transcribed and this process strongly impacts gene regulation and chromatin structure (60, 61). In prokaryotes, various potential roles of pervasive antisense transcription in gene regulation and genome evolution were considered (62) but, to date, such roles have remained hypothetical. Data presented here show that the elongation of pervasive transcripts into H-NS-DNA complexes can act as a counter-silencing mechanism modulating a regulatory response. Although most of the effects were observed under conditions of impaired transcription termination, low-level readthrough transcripts were detected in unchallenged cells suggesting that their effects (*e.g*., bistability) are exerted during normal growth. This study adds elongation of pervasive transcripts to the set of mechanisms that produce transcriptional noise (63) and provides a model to understand the molecular basis of SPI-1 bistability, which has remained a longstanding mystery in *Salmonella* biology. The model fits well in the view that stochastic cell-to-cell differences perpetuated by feedback loops can generate phenotypic lineages (64, 65).

## Materials and Methods

### Strains and culture conditions

All strains used in this work are derived from *Salmonella enterica* serovar Typhimurium strain LT2 (66). Strains and their genotypes are listed in Supplementary Table S1. Bacteria were routinely cultured in Lysogeny Broth (LB: Tryptone 10 g/l, Yeast extract 5 g/l, NaCl, 5 g/l) at 37°C or, occasionally, at 30°C when carrying temperature-sensitive plasmid replicons. Typically, bacteria were grown overnight in static 2 ml cultures (14 mm diameter tubes), subcultured by 1:200 dilution the next day (20 ml culture in 125 ml Erlenmeyer flasks) and grown with 170 rpm shaking. For growth on plates, LB was solidified by the addition of 1.5% Difco agar. When needed, antibiotics (Sigma-Aldrich) were included in growth media at the following final concentrations: chloramphenicol, 10 μg/ml; kanamycin monosulphate, 50 μg/ml; sodium ampicillin 100 μg/ml; spectinomycin dihydrochloride, 80 μg/ml; tetracycline hydrochloride, 25 μg/ml. Strains were constructed by generalized transduction using the high-frequency transducing mutant of phage P22, HT 105/1 *int-201* (67) or by the *l-red* recombineering technique implemented previously (68). 3xFLAG epitope fusions were constructed as described (69) or by two-step scarless recombineering. The latter procedure involved the use of tripartite selectable counter-selectable cassettes (conditionally expressing the *ccdB* toxin gene) amplified from in-house-developed plasmid templates. Oligonucleotide used as primers for amplification (obtained from Sigma-Aldrich or Eurofins) are listed in Supplementary Table S2. Their assortment for the construction of the relevant alleles used in this study is shown in Supplementary Table S3. PCR-amplified fragments to be used for recombineering were produced with high-fidelity Phusion polymerase (New England Biolabs). Constructs were verified by colony-PCR using Taq polymerase followed by DNA sequencing (performed by Eurofins-GATC Biotech).

### Fluorescence microscopy

Bacterial cultures grown overnight in LB at 37°C were diluted 1:200 into 2 ml of the same medium with or without 0.1% arabinose and/or 0.4 μg/ml AHTc (in 14 mm diameter tubes) and grown for 4 hours at 37°C with shaking (170 rpm). Cells were then harvested by centrifugation (2 min at 12,000 x g), washed once in PBS and used immediately for microscopic examination. Images were captured with a Leica DM 6000 B microscope (CTR 6500 drive control unit) equipped with a EBQ 100 lamp power unit and filters for phase contrast, GFP and mCherry detection (100 x oil immersion objective). Pictures were taken with a Hamamatsu C11440 digital camera and processed with Metamorph software.

### Flow cytometry

Flow cytometry was used to monitor expression of translational GFP fusions. Data acquisition was performed using a Cytomics FC500-MPL cytometer (Beckman Coulter, Brea, CA) and data were analyzed with FlowJo X version 10.0.7r software (Tree Star, Inc., Ashland, OR). *S. enterica* cultures were washed and re-suspended in phosphate-buffered saline (PBS) for fluorescence measurement. Fluorescence values for 100,000 events were compared with the data from the reporterless control strain, thus yielding the fraction of ON and OFF cells.

### RNA extraction and quantification by RT-qPCR

Overnight bacterial cultures in LB were diluted 1:200 in the same medium – or in LB supplemented with 0.1% ARA or 0.4 μg/ml Anhydrotetracycline (AHTc) or both drugs where appropriate – and grown with shaking at 37°C to an OD_600_ = 0.7 to 0.8. Cultures (4 ml) were rapidly spun down and resuspended in 0.6 ml ice-cold REB buffer (20 mM Sodium Acetate pH 5.0, 10% sucrose). RNA was purified by sequential extraction with hot acid phenol, phenolchloroform 1:1 mixture and chloroform. Following overnight ethanol precipitation at −20°C and centrifugation, the RNA pellet was resuspended in 20 μl of H_2_O. Three samples were prepared from independent biological replicates for each strain and condition. RNA yields, measured by Nanodrop reading, typically ranged between 2 and 3 μg/μl. The RNA preparations were used for first-strand DNA synthesis with the New England Biolabs (NEB) ProtoScript II First Strand DNA synthesis kit, following the manufacturer’s specifications. Briefly, RNA (1 μg) was combined with 2 μl of a mixture of two primers (5 μM each), one annealing in the promoter proximal portion of the RNA to be quantified (primer AI41 for *hilD* or primer AI48 for *prgH*), the other annealing to a similar position in the reference RNA (primer AJ33 for *ompA*) in an 8 μl final volume. After 5’ min at 65°C and a quick cooling step on ice, volumes were brought to 20 μl by the addition of 10 μl of ProtoScript II Reaction Mix (2x) and 2 μl of ProtoScript II Enzyme Mix (10x). Mixes were incubated for one hour at 42°C followed by a 5 min enzyme inactivation step at 80°C. Samples were then used for real time quantitative PCR as described in Supplementary Materials and Methods.

### 5’RACE-Seq analysis

RNA 5’-end analysis was carried out by template switching reverse transcription (51) coupled to PCR. Initially, we applied this technique on RNA pretreated with Vaccinia virus capping enzyme as reported previously (70). However, these initial tests indicated that the capping step is unnecessary; therefore, this step was subsequently omitted. From that point on, we followed the protocol described by the Template Switching RT Enzyme Mix provider (New England Biolabs) with a few modifications (Supplementary Materials and Methods). The synthesized cDNA was amplified by PCR with primers carrying Illumina adapters at their 5’ ends. Several PCRs were carried out in parallel with a common forward primer (AJ38, annealing to the TSO) and a reverse primer specific for the region being analysed and carrying a treatment-specific index sequence (see example in Supplementary Fig. S3). Reactions were set up according to New England Biolabs PCR protocol for Q5 Hot Start High-Fidelity DNA polymerase in a final volume of 50 μl (using 1 μl of the above cDNA preparation per reaction). The number of amplification cycles needed for reproducible semiquantitative measurements, determined in trial experiments, was chosen to be 25 for the *ompA* reference, 30 for the primary *hilD* and *prgH* promoters and 35 for the secondary *hilD* promoters and the P^tetA^ promoter. The PCR program was as follows: activation: 98°C for 30 sec; amplification (25 or 30 or 35 cycles): 98°C for 10 sec; 65°C for 15 sec; 72°C for 30 sec; final stage: 72°C for 5 min. Products from parallel PCRs were mixed in equal volumes; mixes originating from the amplification of separate regions were pooled and the pools subjected to high throughput sequencing. The procedure was implemented at least once, occasionally twice, with each of the independent RNA preparations. The counts of reads containing the TSO sequence positioned at the TSSs analysed here, each normalized to the counts of reads containing the TSO positioned at the *ompA* TSS, were used to calculate the ratios between the activity of a promoter under a given treatment relative and its activity in untreated cells. The raw data from RACE-Seq experiments were deposited into ArrayExpress under the accession number E-MTAB-11419.

### ChIP-Seq analysis

Overnight bacterial cultures were diluted 1:100 in LB or in LB supplemented with 0.1% ARA or 0.1% ARA + 0.4 μg/ml AHTc and grown at 37°C to an OD600 of 0.7-0.8. At this point 1.6 ml of 37% Formaldehyde (Alfa Aesar) were added to 30 ml of culture and the culture incubated for 30 min at room temperature with gentle agitation. This was followed by the addition of 6.8 ml of a 2.5 M glycine solution and further 15 min incubation with gentle agitation at room temperature. Cells were centrifuged and the pellet resuspended in 24 ml of TBS buffer (50 mM Tris HCl pH7.4, 150mM NaCl). These steps were repeated once and the cells centrifuged again. Cells were then processed for ChIP as previously described (71) and adapted here to *Salmonella* (see Supplementary Materials and Methods). The raw data from all ChIP-Seq experiments were deposited into ArrayExpress under the accession number E-MTAB-11386.

## Statistics, Reproducibility and Bioinformatic analyses

See Supplementary Materials and Methods.

## Acknowledgments

We are grateful to Vicky Lioy for advice on ChIP experiments and to Modesto Carballo, Laura Navarro and Cristina Reyes (Servicio de Biología, CITIUS, Universidad de Sevilla) for help with flow cytometry analysis. We thank the High-throughput Sequencing Core Facility of the I2BC (Gif-sur-Yvette) for library preparation and sequencing (ChIP-Seq and RACE-Seq) and the ICGex NGS platform of the Curie Institute (Paris) for generating some of the sequence data sets (RACE-Seq). This study was supported by the Centre National de la Recherche Scientifique (CNRS), by the Agence Nationale de la Recherche (ANR-15-CE11-0024-03) France and by Grant BIO2016-75235-P (Ministerio de Ciencia e Innovación, Spain and the European Regional Fund). ChIP-Seq data and RACE-Seq data were deposited into ArrayExpress under the accession numbers E-MTAB-11386 and E-MTAB-11419, respectively.

## References

1. J. M. Peters et al., Rho and NusG suppress pervasive antisense transcription in Escherichia coli. Genes Dev 26, 2621–2633 (2012).

2. J. M. Peters et al., Rho directs widespread termination of intragenic and stable RNA transcription. Proc Natl Acad Sci U S A 106, 15406–15411 (2009).

3. L. Ettwiller, J. Buswell, E. Yigit, I. Schildkraut, A novel enrichment strategy reveals unprecedented number of novel transcription start sites at single base resolution in a model prokaryote and the gut microbiome. BMC Genomics 17, 199 (2016).

4. V. V. Panyukov, O. N. Ozoline, Promoters of Escherichia coli versus promoter islands: function and structure comparison. PLoS One 8, e62601 (2013).

5. S. S. Singh et al., Widespread suppression of intragenic transcription initiation by H-NS. Genes Dev 28, 214–219 (2014).

6. D. Forrest, E. A. Warman, A. M. Erkelens, R. T. Dame, D. C. Grainger, Xenogeneic silencing strategies in bacteria are dictated by RNA polymerase promiscuity. Nat Commun 13, 1149 (2022).

7. L. E. Lamberte et al., Horizontally acquired AT-rich genes in Escherichia coli cause toxicity by sequestering RNA polymerase. Nat Microbiol 2, 16249 (2017).

8. R. Landick, J. T. Wade, D. C. Grainger, H-NS and RNA polymerase: a love-hate relationship? Curr Opin Microbiol 24, 53–59 (2015).

9. C. J. Dorman, H-NS: a universal regulator for a dynamic genome. Nat Rev Microbiol 2, 391-400 (2004).

10. C. J. Dorman, J. C. Hinton, A. Free, Domain organization and oligomerization among H-NS-like nucleoid-associated proteins in bacteria. Trends Microbiol 7, 124–128 (1999).

11. F. C. Fang, S. Rimsky, New insights into transcriptional regulation by H-NS. Curr Opin Microbiol 11, 113–120 (2008).

12. S. T. Arold, P. G. Leonard, G. N. Parkinson, J. E. Ladbury, H-NS forms a superhelical protein scaffold for DNA condensation. Proc Natl Acad Sci U S A 107, 15728–15732 (2010).

13. C. Ueguchi, T. Mizuno, The Escherichia coli nucleoid protein H-NS functions directly as a transcriptional repressor. Embo j 12, 1039–1046 (1993).

14. R. T. Dame, C. Wyman, R. Wurm, R. Wagner, N. Goosen, Structural basis for H-NS-mediated trapping of RNA polymerase in the open initiation complex at the rrnB P1. J Biol Chem 277, 2146–2150 (2002).

15. M. Shin et al., DNA looping-mediated repression by histone-like protein H-NS: specific requirement of Esigma70 as a cofactor for looping. Genes Dev 19, 2388–2398 (2005).

16. L. Bossi et al., NusG prevents transcriptional invasion of H-NS-silenced genes. PLoS Genet 15, e1008425 (2019).

17. M. V. Kotlajich et al., Bridged filaments of histone-like nucleoid structuring protein pause RNA polymerase and aid termination in bacteria. Elife 4 (2015).

18. S. Lucchini et al., H-NS mediates the silencing of laterally acquired genes in bacteria. PLoS Pathog 2, e81 (2006).

19. W. W. Navarre et al., Selective silencing of foreign DNA with low GC content by the H-NS protein in Salmonella. Science 313, 236–238 (2006).

20. S. S. Ali, B. Xia, J. Liu, W. W. Navarre, Silencing of foreign DNA in bacteria. Curr Opin Microbiol 15, 175–181 (2012).

21. C. J. Dorman, H-NS, the genome sentinel. Nat Rev Microbiol 5, 157–161 (2007).

22. S. S. Ali et al., Silencing by H-NS potentiated the evolution of Salmonella. PLoS Pathog 10, e1004500 (2014).

23. K. Higashi et al., H-NS Facilitates Sequence Diversification of Horizontally Transferred DNAs during Their Integration in Host Chromosomes. PLoS Genet 12, e1005796 (2016).

24. J. C. Perez, T. Latifi, E. A. Groisman, Overcoming H-NS-mediated transcriptional silencing of horizontally acquired genes by the PhoP and SlyA proteins in Salmonella enterica. J Biol Chem 283, 10773–10783 (2008).

25. D. M. Stoebel, A. Free, C. J. Dorman, Anti-silencing: overcoming H-NS-mediated repression of transcription in Gram-negative enteric bacteria. Microbiology (Reading, England) 154, 2533–2545 (2008).

26. W. R. Will, D. H. Bale, P. J. Reid, S. J. Libby, F. C. Fang, Evolutionary expansion of a regulatory network by counter-silencing. Nat Commun 5, 5270 (2014).

27. W. R. Will, W. W. Navarre, F. C. Fang, Integrated circuits: how transcriptional silencing and counter-silencing facilitate bacterial evolution. Curr Opin Microbiol 23, 8–13 (2015).

28. J. R. Ellermeier, J. M. Slauch, Adaptation to the host environment: regulation of the SPI1 type III secretion system in Salmonella enterica serovar Typhimurium. Curr Opin Microbiol 10, 24–29 (2007).

29. J. E. Galán, Salmonella interactions with host cells: type III secretion at work. Annual review of cell and developmental biology 17, 53–86 (2001).

30. C. D. Ellermeier, J. R. Ellermeier, J. M. Slauch, HilD, HilC and RtsA constitute a feed forward loop that controls expression of the SPI1 type three secretion system regulator hilA in Salmonella enterica serovar Typhimurium. Mol Microbiol 57, 691–705 (2005).

31. J. Lopez-Garrido, E. Puerta-Fernandez, J. Casadesus, A eukaryotic-like 3’ untranslated region in Salmonella enterica hilD mRNA. Nucleic Acids Res 42, 5894–5906 (2014).

32. C. C. Hung et al., Salmonella invasion is controlled through the secondary structure of the hilD transcript. PLoS Pathog 15, e1007700 (2019).

33. D. Pérez-Morales et al., An incoherent feedforward loop formed by SirA/BarA, HilE and HilD is involved in controlling the growth cost of virulence factor expression by Salmonella Typhimurium. PLoS Pathog 17, e1009630 (2021).

34. J. R. Grenz, J. E. Cott Chubiz, P. Thaprawat, J. M. Slauch, HilE Regulates HilD by Blocking DNA Binding in Salmonella enterica Serovar Typhimurium. J Bacteriol 200, (2018).

35. I. N. Olekhnovich, R. J. Kadner, DNA-binding activities of the HilC and HilD virulence regulatory proteins of Salmonella enterica serovar Typhimurium. J Bacteriol 184, 4148–4160 (2002).

36. C. Smith, A. M. Stringer, C. Mao, M. J. Palumbo, J. T. Wade, Mapping the Regulatory Network for Salmonella enterica Serovar Typhimurium Invasion. MBio 7 (2016).

37. M. Kalafatis, J. M. Slauch, Long-Distance Effects of H-NS Binding in the Control of hilD Expression in the Salmonella SPI1 Locus. J Bacteriol 203, e0030821 (2021).

38. M. Ackermann et al., Self-destructive cooperation mediated by phenotypic noise. Nature 454, 987–990 (2008).

39. M. Diard et al., Stabilization of cooperative virulence by the expression of an avirulent phenotype. Nature 494, 353–356 (2013).

40. I. Hautefort, M. J. Proença, J. C. Hinton, Single-copy green fluorescent protein gene fusions allow accurate measurement of Salmonella gene expression in vitro and during infection of mammalian cells. Appl Environ Microbiol 69, 7480–7491 (2003).

41. S. Saini, J. R. Ellermeier, J. M. Slauch, C. V. Rao, The role of coupled positive feedback in the expression of the SPI1 type three secretion system in Salmonella. PLoS Pathog 6, e1001025 (2010).

42. M. A. Sánchez-Romero, J. Casadesús, Contribution of SPI-1 bistability to Salmonella enterica cooperative virulence: insights from single cell analysis. Scientific reports 8, 14875 (2018).

43. M. C. Schlumberger et al., Real-time imaging of type III secretion: Salmonella SipA injection into host cells. Proc Natl Acad Sci U S A 102, 12548–12553 (2005).

44. M. A. Sánchez-Romero, J. Casadesús, Single Cell Analysis of Bistable Expression of Pathogenicity Island 1 and the Flagellar Regulon in Salmonella enterica. Microorganisms 9 (2021).

45. A. Sturm et al., The cost of virulence: retarded growth of Salmonella Typhimurium cells expressing type III secretion system 1. PLoS Pathog 7, e1002143 (2011).

46. M. Arnoldini et al., Bistable expression of virulence genes in salmonella leads to the formation of an antibiotic-tolerant subpopulation. PLoS Biol 12, e1001928 (2014).

47. B. Stecher et al., Salmonella enterica serovar typhimurium exploits inflammation to compete with the intestinal microbiota. PLoS Biol 5, 2177–2189 (2007).

48. L. Bossi, A. Schwartz, B. Guillemardet, M. Boudvillain, N. Figueroa-Bossi, A role for Rho-dependent polarity in gene regulation by a noncoding small RNA. Genes Dev 26, 1864–1873 (2012).

49. C. P. Lostroh, C. A. Lee, The HilA box and sequences outside it determine the magnitude of HilA-dependent activation of P(prgH) from Salmonella pathogenicity island 1. J Bacteriol 183, 4876–4885 (2001).

50. R. A. Daly, C. P. Lostroh, Genetic analysis of the Salmonella transcription factor HilA. Canadian journal of microbiology 54, 854–860 (2008).

51. M. G. Wulf et al., Non-templated addition and template switching by Moloney murine leukemia virus (MMLV)-based reverse transcriptases co-occur and compete with each other. J Biol Chem 294, 18220–18231 (2019).

52. C. Kröger et al., An infection-relevant transcriptomic compendium for Salmonella enterica Serovar Typhimurium. Cell Host Microbe 14, 683–695 (2013).

53. N. A. Lerminiaux, K. D. MacKenzie, A. D. S. Cameron, Salmonella Pathogenicity Island 1 (SPI-1): The Evolution and Stabilization of a Core Genomic Type Three Secretion System. Microorganisms 8 (2020).

54. E. Bouffartigues, M. Buckle, C. Badaut, A. Travers, S. Rimsky, H-NS cooperative binding to high-affinity sites in a regulatory element results in transcriptional silencing. Nat Struct Mol Biol 14, 441–448 (2007).

55. V. H. Bustamante et al., HilD-mediated transcriptional cross-talk between SPI-1 and SPI-2. Proc Natl Acad Sci U S A 105, 14591–14596 (2008).

56. L. C. Martínez, M. M. Banda, M. Fernández-Mora, F. J. Santana, V. H. Bustamante, HilD induces expression of Salmonella pathogenicity island 2 genes by displacing the global negative regulator H-NS from ssrAB. J Bacteriol 196, 3746–3755 (2014).

57. A. A. Rangarajan, K. Schnetz, Interference of transcription across H-NS binding sites and repression by H-NS. Mol Microbiol 108, 226–239 (2018).

58. J. T. Wade, D. C. Grainger, Waking the neighbours: disruption of H-NS repression by overlapping transcription. Mol Microbiol 108, 221–225 (2018).

59. H. Leh et al., Bacterial-Chromatin Structural Proteins Regulate the Bimodal Expression of the Locus of Enterocyte Effacement (LEE) Pathogenicity Island in Enteropathogenic Escherichia coli. mBio 8 (2017).

60. J. Soudet, F. Stutz, Regulation of Gene Expression and Replication Initiation by Non-Coding Transcription: A Model Based on Reshaping Nucleosome-Depleted Regions: Influence of Pervasive Transcription on Chromatin Structure. Bioessays 41, e1900043 (2019).

61. K. Struhl, Transcriptional noise and the fidelity of initiation by RNA polymerase II. Nat Struct Mol Biol 14, 103–105 (2007).

62. J. T. Wade, D. C. Grainger, Pervasive transcription: illuminating the dark matter of bacterial transcriptomes. Nat Rev Microbiol 12, 647–653 (2014).

63. R. Silva-Rocha, V. de Lorenzo, Noise and robustness in prokaryotic regulatory networks. Annu Rev Microbiol 64, 257–275 (2010).

64. M. Kaern, T. C. Elston, W. J. Blake, J. J. Collins, Stochasticity in gene expression: from theories to phenotypes. Nat Rev Genet 6, 451–464 (2005).

65. M. A. Sanchez-Romero, J. Casadesus, Waddington’s Landscapes in the Bacterial World. Front Microbiol 12, 685080 (2021).

66. K. Lilleengen, Typing of *Salmonella typhimurium* by means of bacteriophage. Acta Pathol. Microbiol. Scand. 77, 2–125 (1948).

67. H. Schmieger, Phage P22-mutants with increased or decreased transduction abilities. Mol Gen Genet 119, 75–88 (1972).

68. K. A. Datsenko, B. L. Wanner, One-step inactivation of chromosomal genes in Escherichia coli K-12 using PCR products. Proc Natl Acad Sci U S A 97, 6640–6645 (2000).

69. S. Uzzau, N. Figueroa-Bossi, S. Rubino, L. Bossi, Epitope tagging of chromosomal genes in Salmonella. Proc Natl Acad Sci U S A 98, 15264–15269 (2001).

70. F. Liu, K. Zheng, H. C. Chen, Z. F. Liu, Capping-RACE: a simple, accurate, and sensitive 5’ RACE method for use in prokaryotes. Nucleic Acids Res 46, e129 (2018).

71. V. S. Lioy, F. Boccard, Conformational Studies of Bacterial Chromosomes by High-Throughput Sequencing Methods. Methods Enzymol 612, 25–45 (2018).

