## Supplementary Materials and Methods plus 6 Supplementary figures and 4 Supplementary tables for "Pervasive transcription enhances the accessibility of H-NS-silenced promoters and generates bistability in *Salmonella* virulence gene expression" (from another version of this preprint: version 1, 2022-04-26)

**This PDF file includes:**
Supplementary text (Materials and Methods)
Figures S1 to S6
Table S1 to S4

**Supplementary Materials and Methods**

***RT-qPCR.*** Amplification reactions were set up in 384-well plates by mixing serial dilutions of each reverse-transcribed sample with the appropriate primer pairs (each primer used at a 0.25 µM final concentration) and the LightCycler 480 SYBR Green I Master Mix (Roche Applied Science). Real-time qPCR was carried out in a LightCycler 480 Instrument (Roche) with the following program: activation: 95°C for 5 min; amplification (40 cycles): 95°C for 10 sec; 55°C for 20 sec; 72°C for 20 sec; melting curve: 95°C for 30 sec; 65°C for 30 sec (ramp 0.06 °C/sec, 10 acquisitions/°C). Target-to-reference transcript ratios and relative transcript levels were calculated with the Pflaffl method (1).

***Template switching Reverse Transcription.*** Total RNA (1 µg) was combined with 1 µl of a mixture of up to four gene-specific primers, 5 µM each (including AI41 (*hilD*), AI48 (*prgH*) and AJ33 (*ompA*)), and 1 µl of 10 mM dNTPs in a 6 µl final volume. After a 5 min treatment at 70°C (in a Thermocycler), samples were quickly cooled on ice. Each sample were then mixed with 2.5 µl of Template Switching Buffer (4x), 0.5 µl of 75 µl of Template Switching Oligonucleotide (TSO) and 1 µl of Template Switching RT Enzyme Mix in a final volume of 10 µl. Reverse transcription was carried out for 90 min at 42°C, followed by a 5 min incubation at 85°C. The 42 nucleotide-long TSO carries three 3'-terminal riboguanosines that hybridize to the polycytosine overhang (typically 3 or 4 Cs) added by the RT enzyme when it reaches the 5' end of the RNA. This allows the RT enzyme to switch template and copy the TSO.

***CHIP-Seq analysis.*** Following two washes with TBS buffer**,** Formaldehyde-treated cells were resuspended in 0.5 ml of Lysis buffer (50mM Tris pH 7.4, 150mM NaCl, 1mM EDTA supplemented with cOmplete™ Mini Protease Inhibitor cocktail (Sigma-Aldrich). Then, 3 µl of Ready-Lyse™ Lysozyme solution, 25,000 – 40,000 U/µl (Lucigen R1804M) were added and incubated on ice for 10 min, followed by the addition of 25 µl of 25% Triton and 450 µl of Lysis buffer. Cells were disrupted in a Covaris 220 focused ultrasonicator (Peak Incidence = 140W; Duty Factor = 5; Cycles per burst = 200). This step was followed by 10 min centrifugation at 16 000 rcf at 4°C. After removing aliquots for "Input" controls, supernatants (0.9 ml) were treated overnight with Anti-Flag M2 affinity Gel (Sigma A2220, 40 µl of slurry per sample) on a gyratory wheel at 8°C. Next day, beads were washed in the cold with 500 µl of TBS plus 0.05% Tween 20 for 10 minutes, followed by four TBS washes, of 10 minute each. The elution of 3xFLAG-H-NS DNA complex from the beads was accomplished as follows: 3xFLAG peptide solution (prepared according to Sigma Aldrich instructions) was diluted with TBS (3 µl of 3xFLAG peptide concentrated solution per 100 µl of TBS) and an aliquot of 2.5 times volume (relative to beads) of the peptide-TBS solution was added on top of the beads. Tubes were incubated for 1 hour on the gyratory wheel at 8°C. Samples were centrifuged briefly and supernatant transferred to a clean tube. Inputs as well as samples were treated with Proteinase K (2.5 µl of a 2 mg/ml solution) at 65°C overnight. DNA was cleaned using Qiagen Mini-elute reaction clean up kit. DNA fragments were end-repaired and dA-tailed (NEB#E7595), Illumina TruSeq adapters were ligated (NEB#E6040) and libraries were amplified with Kapa Hifi polymerase (Kapa Biosystem #KK2103). Final libraries quality was assessed on an Agilent Bioanalyzer 2100, using an Agilent High Sensitivity DNA Kit.

Libraries were pooled in equimolar proportions and sequenced on a Paired-End 2x75 bp run, on an Illumina NextSeq500 instrument. Output files were demultiplexed and adapters were removed. Only reads longer than 10 pb were kept for analysis. The raw data from all CHIP-Seq experiments were deposited into ArrayExpress under the accession number E-MTAB-11386.

***Measurement of ß-galactosidase activity.*** Static overnight cultures were diluted 1:200 in 2 ml of LB (with or without ARA) and grown with shaking at 37°C to early or mid-stationary phase. Bacteria were harvested from 1 ml aliquots and resuspended in 1 ml of PBS. β-galactosidase activity was assayed in toluene-permeabilized cells as described by Miller (2) and is expressed in Miller units. Measurements were repeated three times, each time using duplicate cultures originating from independent colonies.

***Statistics and Reproducibility.*** All data described in this paper originate from three or more independent experiments, with one of more measurements performed on each replicate of the experiment. Statistical significance was calculated as specified in the legends to the figures. In pairwise comparisons of expression levels, significance was generally determined by unpaired two-tailed Student's t-tests with Welch's correction for unequal variances. In flow cytometry experiments, significance was determined performing one-way ANOVA with Dunnett's multiple comparisons test. All statistical analyses were done GraphPad Prism 9 software. P values were included in the figures (using the asterisk symbol) or specified in the figure legends.

***Bioinformatic analyses.*** Demultiplexing of raw data from the Illumina sequencer was performed with the bcl2fastq2 V2.2.18.12 program and adapters were trimmed with Cutadapt1.15. The reads from the CHIP-Seq experiments were mapped on the genomes of *Salmonella enterica* serovar Typhimurium strain LT2 (wt) and the SPI-1-modified variant MA14358 with BWA 0.6.2-r126. Bedgraph files were generated from aligned Bam files using bedtools genomecov. Coverage track (number of reads per base) was converted to the BigWig format using the bedGraphToBigWig command line utility from UCSC. Read depth was calculated using the bedcov tool of the Samtools suite. Bam and Bigwig files were visualized with Integrative Genome Viewer (IGV) (3). In the processing of the RACE-Seq data, the reads containing the TSO were filtered and kept thanks to grep command in the SeqKit package. These reads were then trimmed with the PRINSEQ tool to remove 33 bp from the 5' end (the amplified portion of the TSO except the terminal 3Gs).

**References**

1. M. W. Pfaffl, A new mathematical model for relative quantification in real-time RT-PCR. *Nucleic Acids Res* **29**, e45 (2001).
2. J. H. Miller, *A Short Course in Bacterial Genetics. A Laboratory Manual and Handbook for Escherichia coli and Related Bacteria.* (Cold Spring Harbor Laboratory Press, Cold Spring Harbor, New York, 1992).
3. J. T. Robinson *et al.*, Integrative genomics viewer. *Nat Biotechnol* **29**, 24-26 (2011).

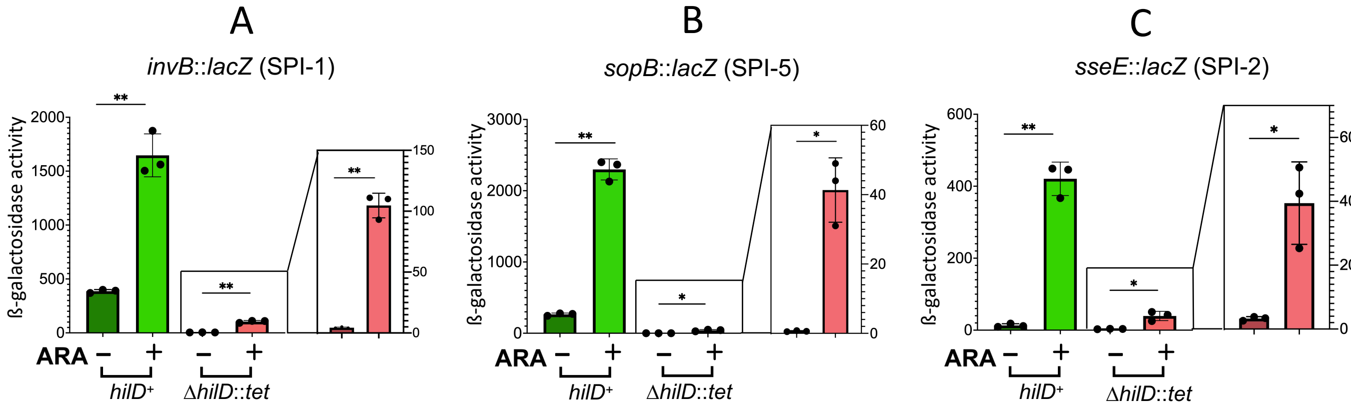


**Figure S1.** Upregulation of SPI genes in NusG-depleted cells is largely HilD-mediated. Strains carrying translational *lacZ* gene fusions to SPI genes in the ARA-inducible NusG depletion background were grown at 37°C in LB (with or without ARA) for 4.5 hours (strains with *invB-lacZ* and *sopB-lacZ*) or for 6 hours (strain with *sseE-lacZ*). At this time, $OD_{600}$ readings ranged between 1.1 and 2.1 (4.5 h) or between 1.9 and 3.3 (6 h) depending on the strain genotype and the presence or absence of ARA. Cells were assayed for ß-galactosidase activity as described in Methods. Activity is expressed in Miller units. (*A*) Strains MA13397 (*hilD*+) and MA13971 (Δ*hilD*::*tet*) carrying *invB-lacZ* (SPI-1). (*B*) Strains MA13398 (*hilD*+) and MA13972 (Δ*hilD*::*tet*) carrying *sopB-lacZ* (SPI-5). (*C*) Strains MA13409 (*hilD*+) and MA13973 (Δ*hilD*::*tet*) carrying *sseE-lacZ* (SPI-2). The results shown represent the means of three independent determinations (with error bars indicating SD) each carried out in duplicate. Significance was determined by unpaired two-tailed Student's t-tests with Welch's correction for unequal variances (*, $P \leq 0.05$; **, $P \leq 0.01$).

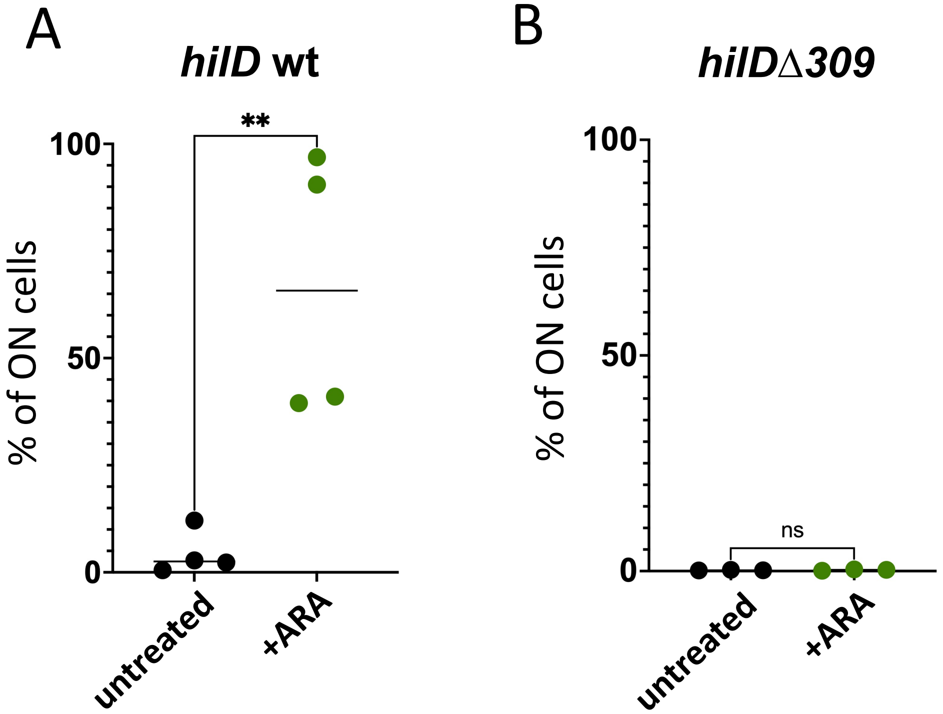


**Figure S2.** NusG depletion enhances HilD-dependent SPI-1 bistability. Flow cytometry analysis of strains MA14302 (*hilD*+) (*A*) and MA14561 (*hilDΔ309*) (*B*). Cells were grown at 37°C in LB with or without ARA added to an $OD_{600}$ of ~0.8. GFP fluorescence intensity was measured, the number of cells analysed as 100,000, and the sizes of the SPI-1$^{ON}$ subpopulations are shown as percentages. The plot represents the results of four (*A*) and three (*B*) independent experiments. Statistical significance was determined by the unpaired two-tailed Student's t-test (ns, $P > 0.05$; **, $P \leq 0.01$).

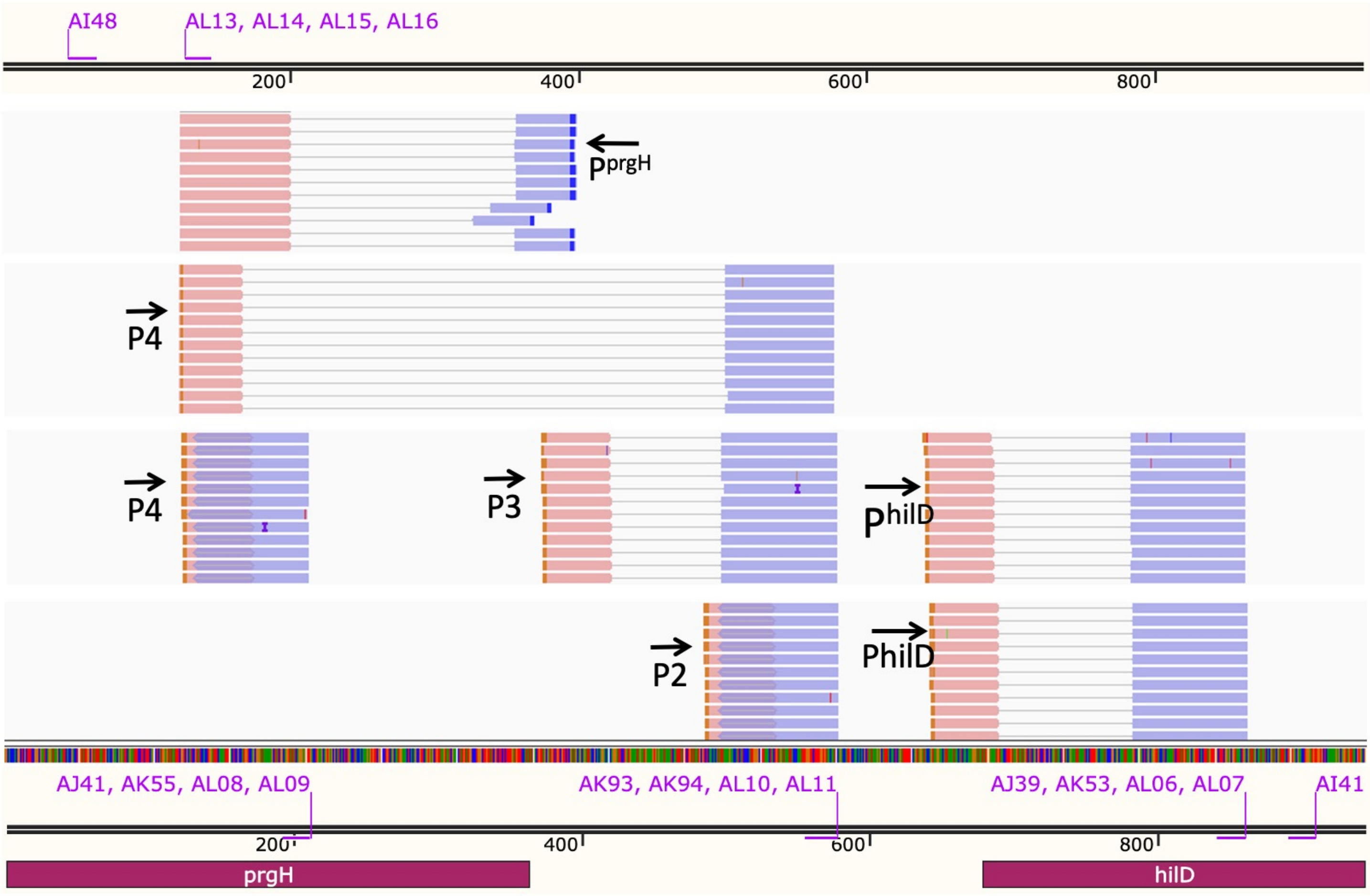


**Figure S3.** Schematic representation of the primer combinations used for 5' RACE Seq. The positions of RT primers (AI41 and AI48) and reverse PCR primers are shown relative to representative reads generated by high-throughput sequencing (Bam files visualized with Integrative genome viewer, IGV).

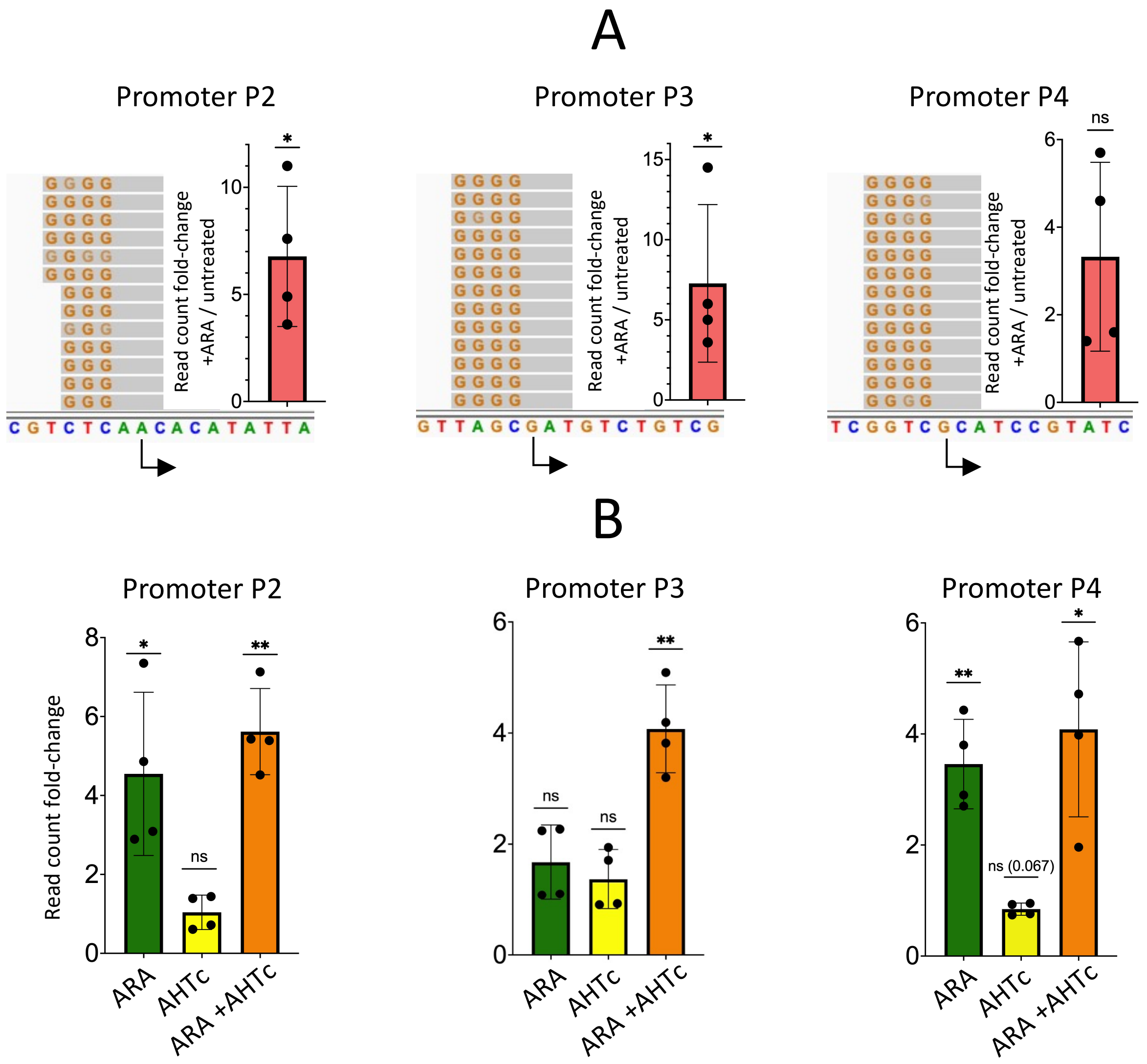


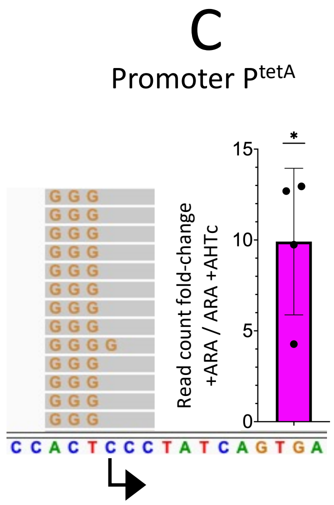


**Figure S4**. 5' RACE-Seq analysis of *hilD* secondary promoters and the $P^{tetA}$ promoter. RNA from strains MA14302 (*A*) and MA14358 (*B* and *C*) was reverse-transcribed in the presence of a template-switching oligonucleotide (TSO). The resulting cDNA was subjected to semi-quantitative PCR amplification (primers shown in Supplementary Fig. S3) and high-throughput sequencing. Read counts were normalized to those measured at the *ompA* promoter. The results shown represent the ratios between the normalized counts from treated and untreated cells. Results originate from three or four independent experiments (with error bars representing SD). Significance was determined by unpaired two-tailed Student's t tests with Welch's correction (ns, P> 0.05; *, P ≤ 0.05; **, P ≤ 0.01; ***, P ≤ 0.001).

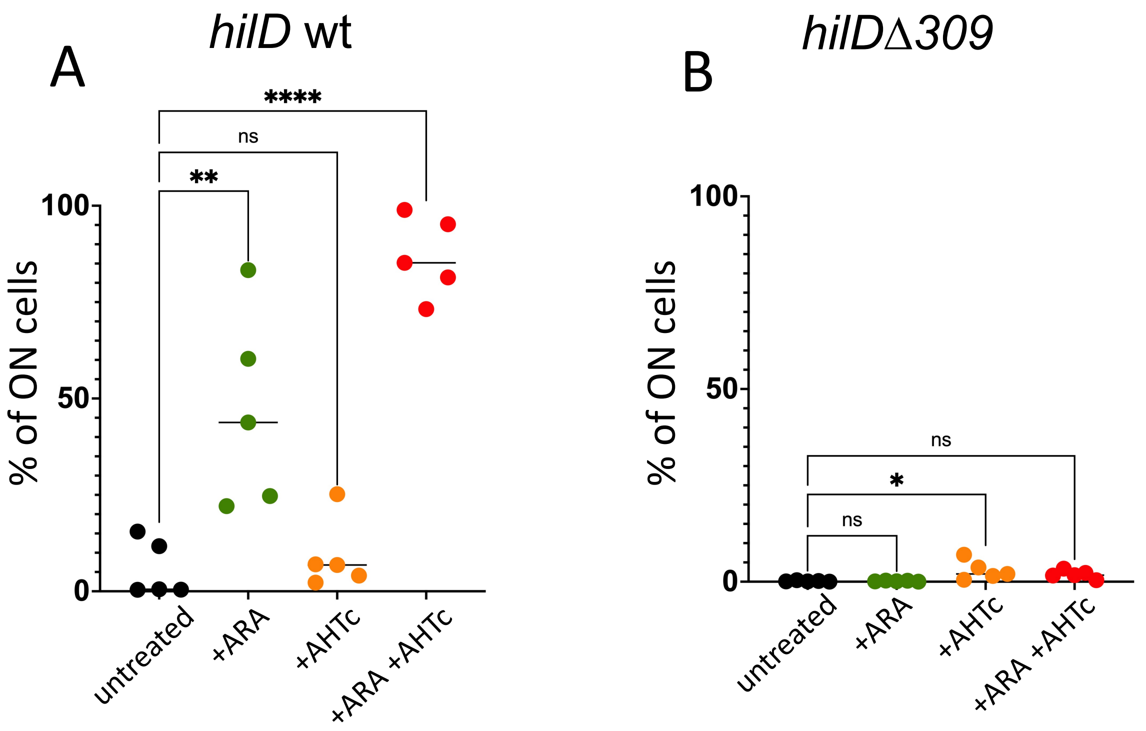


**Figure S5.** Overlapping transcription promotes HilD-dependent bistability of SPI-1 expression. Flow cytometry analysis of strains MA14358 (*tetR*-P$^{tetA}$ *hilD*$^{+}$) (*A*) and MA14569 (*tetR*-P$^{tetA}$ *hilD*Δ*309*) (*B*). Cells were grown at 37°C in plain LB or in LB supplemented with either ARA or AHTc or both. GFP fluorescence intensity was measured, the number of cells analyzed as 100,000 and the sizes of the SPI-1$^{ON}$ subpopulations are shown as percentages. The plot represents the results from five independent experiments. Statistical significance was determined by one-way ANOVA with Dunnett's multiple comparisons test (ns, P> 0.05; *, P ≤ 0.05; ***, P ≤ 0.001; ****, P ≤ 0.0001).

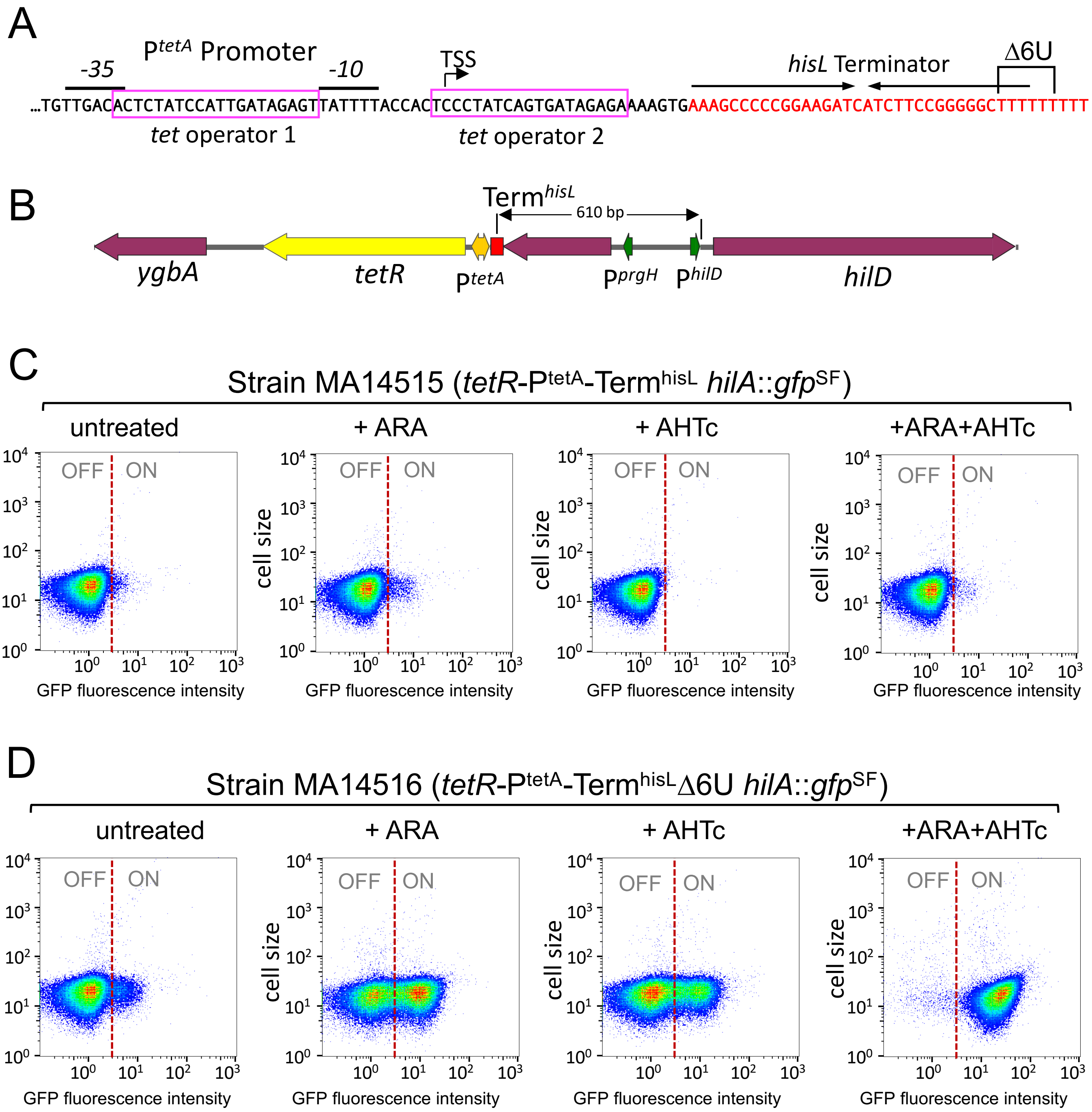


**Figure S6.** Rho-independent transcription terminator abolishes the effects of NusG depletion on SPI-1 bistability. *(A)* Sequence of the P*tetA*-Term*hisL* fusion in strains MA14515 (wt terminator) and MA14516 (Δ6U mutant), respectively. (*B*) Schematic diagram showing Term*hisL* position relative to PtetA and P*hilD* (secondary *hilD* promoters not shown). (*C*) and (*D*) Representative flow cytometry analysis of cells from strains MA14515 (*C*) and MA14516 (*D*). See legend in next page for details.

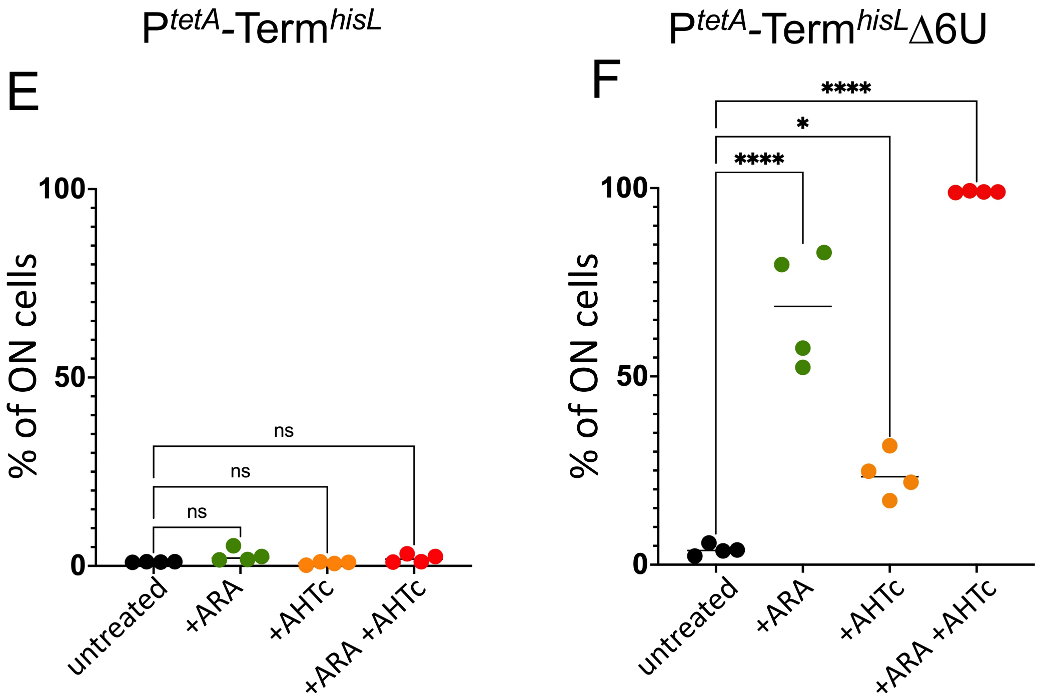


**Figure S6 continued.** Rho-independent transcription terminator abolishes the effects of NusG depletion on SPI-1 bistability. Flow cytometry analysis of strains MA14515 (wt terminator) (*E*) and MA14516 (Δ6U mutant) (*F*), respectively. Cells were grown at 37°C in plain LB or in LB supplemented with either ARA or AHTc or both. GFP fluorescence intensity was measured, the number of cells analyzed as 100,000 and the sizes of the SPI-1$^{ON}$ subpopulations are shown as percentages. The plot represents the results from four independent experiments. Statistical significance was determined by one-way ANOVA with Dunnett's multiple comparisons test (ns, P> 0.05; *, P ≤ 0.05; ***, P ≤ 0.001; ****, P ≤ 0.0001).

### Table S1

**Supplementary Table 1. *Salmonella enterica* serovar Typhimurium strains used in this work**

| Strain | Genotype | Source or reference |
|---|---|---|
| MA3409 | Δ[Gifsy-1] | https://www.ncbi.nlm.nih.gov/pubmed/11902718 |
| MA12603 | Δ[Gifsy-1] hilD-tetR-P^tet-ccdB-cat | this work |
| MA12604 | Δ[Gifsy-1] hilD-3xFLAG | this work |
| MA12996 | Δ[Gifsy-1] Δ[Gifsy-2] Δ[araBAD]::gtgR-P^G2R-nusG-cat Δ[nusG]::aadA | https://pubmed.ncbi.nlm.nih.gov/31589608/ |
| MA13397 | Δ[Gifsy-1] Δ[Gifsy-2] Δ[araBAD]::gtgR-P^G2R-nusG-cat Δ[nusG]::aadA invB-lacZY_kan | https://pubmed.ncbi.nlm.nih.gov/31589608/ |
| MA13398 | Δ[Gifsy-1] Δ[Gifsy-2] Δ[araBAD]::gtgR-P^G2R-nusG-cat Δ[nusG]::aadA sopB-lacZY_kan | https://pubmed.ncbi.nlm.nih.gov/31589608/ |
| MA13409 | Δ[Gifsy-1] Δ[Gifsy-2] Δ[araBAD]::gtgR-P^G2R-nusG-cat Δ[nusG]::aadA sseE-lacZY_kan | https://pubmed.ncbi.nlm.nih.gov/31589608/ |
| MA13748 | Δ[Gifsy-1] Δ[Gifsy-2] Δ[araBAD]::gtgR-P^G2R-nusG-cat Δ[nusG]::aadA hns-3xFLAG_FRT-kan-FRT | this work |
| MA13971 | Δ[Gifsy-1] Δ[Gifsy-2] Δ[araBAD]::gtgR-P^G2R-nusG-cat Δ[nusG]::aadA invB-lacZY_kan ΔhilD::tetRA | this work |
| MA13972 | Δ[Gifsy-1] Δ[Gifsy-2] Δ[araBAD]::gtgR-P^G2R-nusG-cat Δ[nusG]::aadA sopB-lacZY_kan ΔhilD::tetRA | https://pubmed.ncbi.nlm.nih.gov/31589608/ |
| MA13973 | Δ[Gifsy-1] Δ[Gifsy-2] Δ[araBAD]::gtgR-P^G2R-nusG-cat Δ[nusG]::aadA sseE-lacZY_kan ΔhilD::tetRA | https://pubmed.ncbi.nlm.nih.gov/31589608/ |
| MA14067 | Δ[Gifsy-1] Δ[Gifsy-2] Δ[araBAD]::gtgR-P^G2R-nusG-cat Δ[nusG]::aadA hilAΔK28::FRT-kan-FRT | this work |
| MA14082 | Δ[Gifsy-1] Δ[Gifsy-2] Δ[araBAD]::gtgR-P^G2R-nusG-cat Δ[nusG]::aadA hilA::gfpSF ΔK28_kan | this work |
| MA14115 | Δ[Gifsy-1] Δ[Gifsy-2] Δ[araBAD]::gtgR-P^G2R-nusG-cat Δ[nusG]::aadA ΔhilA338::gfpSF | this work |
| MA14122 | Δ[Gifsy-1] Δ[Gifsy-2] Δ[araBAD]::gtgR-P^G2R-nusG-cat Δ[nusG]::aadA ΔinvB1::gfpSF | this work |
| MA14271 | Δ[Gifsy-1] Δ[Gifsy-2] Δ[araBAD]::gtgR-P^G2R-nusG-cat Δ[nusG]::aadA Δ[hisGDCBHAFIE]::P^Tac-mCherry_FRT-kan-FRT | this work |
| MA14275 | Δ[Gifsy-1] Δ[Gifsy-2] Δ[araBAD]::gtgR-P^G2R-nusG-cat Δ[nusG]::aadA Δ[hisGDCBHAFIE]::P^Tac-mCherry_FRT | this work |
| MA14302 | Δ[Gifsy-1] Δ[Gifsy-2] Δ[araBAD]::gtgR-P^G2R-nusG-cat Δ[nusG]::aadA Δ[hisGDCBHAFIE]::P^Tac-mCherry hilA::gfpSF ΔK28_kan | this work |
| MA14315 | Δ[Gifsy-1] Δ[hisGDCBHAFIE]::P^Tac-mCherry hilA::gfpSF ΔK28_kan | this work |
| MA14332 | Δ[Gifsy-1] Δ[hisGDCBHAFIE]::P^Tac-mCherry Δ[sitA-orgA]::tetR-P^tet-ccdB-cat hilA::gfpSF ΔK28_kan | this work |
| MA14339 | Δ[Gifsy-1] Δ[hisGDCBHAFIE]::P^Tac-mCherry Δ[sitA-prgH332]::tetR-P^tet hilA::gfpSF ΔK28_kan | this work |
| MA14345 | Δ[Gifsy-1] Δ[Gifsy-2] Δ[araBAD]::gtgR-P^G2R-nusG-cat Δ[nusG]::aadA orgB::tetR-P^tet-ccdB-kan | this work |
| MA14358 | Δ[Gifsy-1] Δ[Gifsy-2] Δ[araBAD]::gtgR-P^G2R-nusG-cat Δ[nusG]::aadA Δ[hisGDCBHAFIE]::P^Tac-mCherry Δ[sitA-prgH332]::tetR-P^tet hilA::gfpSF ΔK28_kan | this work |
| MA14363 | Δ[Gifsy-1] Δ[Gifsy-2] Δ[araBAD]::gtgR-P^G2R-nusG-cat Δ[nusG]::aadA hilD-3xFLAG | this work |
| MA14505 | Δ[Gifsy-1] Δ[Gifsy-2] Δ[araBAD]::gtgR-P^G2R-nusG-cat Δ[nusG]::aadA Δ[hisGDCBHAFIE]::P^Tac-mCherry Δ[sitA-prgH332]::tetR-P^tet hilD-3xFLAG hilA::gfpSF ΔK28_kan | this work |
| MA14513 | Δ[Gifsy-1] Δ[Gifsy-2] Δ[araBAD]::gtgR-P^G2R-nusG-cat Δ[nusG]::aadA Δ[sitA-prgH332]::tetR-P^tet hilA::gfpSF ΔK28_kan hns-3xFLAG_FRT | this work |
| MA14515 | Δ[Gifsy-1] Δ[Gifsy-2] Δ[araBAD]::gtgR-P^G2R-nusG-cat Δ[nusG]::aadA Δ[hisGDCBHAFIE]::P^Tac-mCherry Δ[sitA-prgH332]::tetR-P^tet-Term^hikL hilA::gfpSF ΔK28_kan | this work |
| MA14516 | Δ[Gifsy-1] Δ[Gifsy-2] Δ[araBAD]::gtgR-P^G2R-nusG-cat Δ[nusG]::aadA Δ[hisGDCBHAFIE]::P^Tac-mCherry Δ[sitA-prgH332]::tetR-P^tet-Term^hikLΔ6U hilA::gfpSF ΔK28_kan | this work |
| MA14558 | Δ[Gifsy-1] Δ[hisGDCBHAFIE]::P^Tac-mCherry hilA::gfpSF ΔK28 hilDΔ309::tetR-P^tet-ccdB-cat | this work |
| MA14560 | Δ[Gifsy-1] Δ[hisGDCBHAFIE]::P^Tac-mCherry hilDΔ309 hilA::gfpSF ΔK28_kan | this work |
| MA14561 | Δ[Gifsy-1] Δ[Gifsy-2] Δ[araBAD]::gtgR-P^G2R-nusG-cat Δ[nusG]::aadA Δ[hisGDCBHAFIE]::P^Tac-mCherry hilDΔ309 hilA::gfpSF ΔK28_kan | this work |
| MA14569 | Δ[Gifsy-1] Δ[Gifsy-2] Δ[araBAD]::gtgR-P^G2R-nusG-cat Δ[nusG]::aadA Δ[hisGDCBHAFIE]::P^Tac-mCherry Δ[sitA-prgH332]::tetR-P^tet hilDΔ309 hilA::gfpSF ΔK28_kan | this work |

### Table S2

**Supplementary Table 2. DNA oligonucleotides used in this work**

| Name | Sequence | Use |
|---|---|---|
| ppY88 | GTGACTGACAATTCATACTATAAATGCGAAATGAAAAAAGCGCTTGACAATTAATCATCGGCTCG | PCR forward primer in λ-red construction (reverse primer is AA30) |
| ppZ13 | TTTAAAACTACGCCATCGACATTCATAAAAATGGCGAACCATTAAGACCCACTTTCACATT | PCR forward primer in λ-red construction |
| ppZ14 | TTTTAATAAAAATCTTTACTTAAGTGACAGATACAAAAAATGCTTACGCCCCGCCCTG | PCR reverse primer in λ-red construction |
| ppZ45 | TTTAAAACTACGCCATCGACATTCATAAAAATGGCGAACCATGACTACAAAGACCATGACGGTGA | PCR forward primer in λ-red construction |
| ppZ46 | TTAATAAAAATCTTTACTTAAGTGACAGATACAAAAAATGTTATTTATCGTCGTCATCTTTGTAGTC | PCR reverse primer in λ-red construction |
| ppW78 | GAAGAACAAGGTAAGCAACTGGAAGATTTCCTGATCAAGGAAGACTACAAAGACCATGACGG | PCR forward primer in λ-red construction |
| ppW79 | AGCGGCGGGATTTTAAGCATCCAGGAAGTAAATTATTCCCATATGAATATCCTCCTTAG | PCR reverse primer in λ-red construction |
| AA30 | CGTAGCTATAAGCCAAAAGCCTTGTAAGCTTCTTTGCCGGATGGTGTAGGCTGGAGCTGCTT | PCR reverse primer in λ-red construction |
| AE12 | GCAAAGGCTATATTCGATGATTAATTAACCACATTGTTGCGAGTTAAGACCCACTTTCACATTTA | PCR forward primer in λ-red construction (reverse primer is AH30) |
| AE45 | GTCATCAGCGTCCTGCCGCAGATAACTTACAGAAATTAAAATCACTTAAGACCCACTTTCACATTTA | PCR forward primer in λ-red construction |
| AE46 | CATTGAATGAAGTAGGACGTGCTATCATAACCACATTTTAATGCCTAAGCACTTGTCTCCTGTTTA | PCR reverse primer in λ-red construction |
| AF19 | GATAGGCGTACATCCGCAAGGTCTTTGCCTCTTCTTCCGCCTCTCTTAAGACCCACTTTCACATTTA | PCR forward primer in λ-red construction (reverse primer is AG26) |
| AF26 | CCGATGGCAAAGCAAAAGTTTAAAATCAACATCGGAGCGGCAGCTGTAGGCTGGAGCTGCTTC | PCR reverse primer in λ-red construction (forward primer is AF93) |
| AF93 | GTACGGACAGGGCTATCGGTTTAATCGTCCGGTCGTAGTGGTGTCGATCCGTCGACCTGCAGTTC | PCR forward primer in λ-red construction |
| AG21 | GTACGGACAGGGCTATCGGTTTAATCGTCCGGTCGTAGTGGTGGCTTCTAAAGGTGAAGAACTGTTC | PCR forward primer in λ-red construction (reverse primer is AG39) |
| AG26 | CTCTATTGAAAGATTAGAACAGCAGGCGCATCAACGGGCTAAGCGGGTCATTGGCATGTTCAA | PCR reverse primer in λ-red construction |
| AG39 | CTGCATCTGAAAAGGAAGTATCGCCAATGTATGAGTCGTAGGTTGCTGTAGGCTGGAGCTGCTTC | PCR reverse primer in λ-red construction |
| AG46 | GTTATAAAAACAATATAAGGGCTTAATTAAGGAAAAGATCTATGGCTTCTAAAGGTGAAGAACTG | PCR forward primer in λ-red construction (reverse primer is AG49) |
| AG49 | GGCCAGATATTGCAGTAAACGAGGTGTTTTCATCTCATTAGCGACCTGTAGGCTGGAGCTGCTTC | PCR reverse primer in λ-red construction |
| AH30 | GGCGGCAAATGAGTTAATACTGGCGGCATGGCGGCTTAAGAACGGTCATTGGCATGTTCAA | PCR reverse primer in λ-red construction |
| AH38 | CATTGATAGAGTTATTTTACCACTCCCTATCAGTGATAGAGAAAAGTGGATAAGCAATTCAC | PCR forward primer in λ-red construction |
| AH39 | CGGTGCAATTAAATACGCCAATACAGGTCGGTGAATTGCTTATCCACTTTTCTC | PCR reverse primer in λ-red construction |
| AI27 | CATTGATAGAGTTATTTTACCACTCCCTATCAGTGATAGAGAAAAGTGAAAGCCCCCGGAAGATCATCTTCCGG | PCR forward primer in λ-red construction |
| AI28 | CGGTGCAATTAAATACGCCAATACAGGTCGGTGAATTGCTTATCAAAAAAAAAGCCCCCGGAAGATGATCTTCCGG | PCR reverse primer in λ-red construction |
| AI39 | GCTAATCATTGCAAGCAGTGGTATCAACGCAGAGTACAT**rGrGrG** | Template-switching oligo (TSO) |
| AI41 | CAACATCCCAGGTTCGTCAC | RT primer (hilD) |
| AI48 | ACCGACCTGTATTGGCGTAT | RT primer (prgH) |
| AI50 | CGCATCCGTATCCACCTGG | qPCR forward primer (prgH) |
| AI51 | GACAGGCCGAACACTCTTTG | qPCR reverse primer (prgH) |
| AI57 | GTGTGTTGGCAATGGTCTGA | qPCR reverse primer (hilD CHIP, forward primer is AL12) |
| AI62 | TGCCGCAGATAACTTACAGAA | qPCR forward primer (hilD) |
| AI63 | GTCAGTTTACCGCTCCGAAA | qPCR reverse primer (hilD) |
| AJ32 | CGTTGGAGATATTCATGGCGT | qPCR forward primer (ompA) |
| AJ33 | ACCAGTCGTAGCCCATTTCA | RT primer (ompA) |
| AJ37 | GAGACCAGCCCAGTTTAGCA | qPCR reverse primer (ompA) |
| AJ38 | AATGATACGGCGACCACCGAGATCTACACTCTTTCCCTACACGACGCTCTTCCGATCTCATTGCAAGCAGTGGTATCAAC | Illumina sequencing universal forward primer (anneals to TSO compleme |
| AJ39 | CAAGCAGAAGACGGCATACGAGAAAGGAATGTGACTGGAGTTCAGACGTGTGCTCTTCCGATCTGTCAGTTTACCGCTCCGAAA | Illumina sequencing reverse primer (hilD primary promoter) |
| AJ40 | CAAGCAGAAGACGGCATACGAGAAAGGAATGTGACTGGAGTTCAGACGTGTGCTCTTCCGATCTTGTGTCCTGTGCGGTAATCT | Illumina sequencing reverse primer (hilD P3 promoter) |
| AJ41 | CAAGCAGAAGACGGCATACGAGAAAGGAATGTGACTGGAGTTCAGACGTGTGCTCTTCCGATCTACTCCCTGATATACCTGCCG | Illumina sequencing reverse primer (hilD P4 promoter and P^tet promote |
| AK19 | GAGTCTTCTGGCTCGCAGTA | qPCR forward primer (katN CHIP) |
| AK20 | CTCTGGCAAGTTTAGGAAGAGC | qPCR reverse primer (katN CHIP) |
| AK53 | CAAGCAGAAGACGGCATACGAGAATAGTTGGTGACTGGAGTTCAGACGTGTGCTCTTCCGATCTGTCAGTTTACCGCTCCGAAA | Illumina sequencing reverse primer (hilD primary promoter) |
| AK54 | CAAGCAGAAGACGGCATACGAGAATAGTTGGTGACTGGAGTTCAGACGTGTGCTCTTCCGATCTTGTGTCCTGTGCGGTAATCT | Illumina sequencing reverse primer (hilD P3 promoter) |
| AK55 | CAAGCAGAAGACGGCATACGAGAATAGTTGGTGACTGGAGTTCAGACGTGTGCTCTTCCGATCTACTCCCTGATATACCTGCCG | Illumina sequencing reverse primer (hilD P4 promoter and P^tet promote |
| AK93 | CAAGCAGAAGACGGCATACGAGAAAGGAATGTGACTGGAGTTCAGACGTGTGCTCTTCCGATCTGATTGTTACACCAAAAGAACCCA | Illumina sequencing reverse primer (hilD P2 and P3 promoters) |
| AK94 | CAAGCAGAAGACGGCATACGAGAATAGTTGGTGACTGGAGTTCAGACGTGTGCTCTTCCGATCTGATTGTTACACCAAAAGAACCCA | Illumina sequencing reverse primer (hilD P2 and P3 promoters) |
| AL01 | CAAGCAGAAGACGGCATACGAGAAAGGAATGTGACTGGAGTTCAGACGTGTGCTCTTCCGATCTGAGACCAGCCCAGTTTAGCA | Illumina sequencing reverse primer (ompA promoter) |
| AL02 | CAAGCAGAAGACGGCATACGAGAATAGTTGGTGACTGGAGTTCAGACGTGTGCTCTTCCGATCTGAGACCAGCCCAGTTTAGCA | Illumina sequencing reverse primer (ompA promoter) |
| AL06 | CAAGCAGAAGACGGCATACGAGAAGTCGTCGTGACTGGAGTTCAGACGTGTGCTCTTCCGATCTGTCAGTTTACCGCTCCGAAA | Illumina sequencing reverse primer (hilD primary promoter) |
| AL07 | CAAGCAGAAGACGGCATACGAGAACTTCGAGTGACTGGAGTTCAGACGTGTGCTCTTCCGATCTGTCAGTTTACCGCTCCGAAA | Illumina sequencing reverse primer (hilD primary promoter) |
| AL08 | CAAGCAGAAGACGGCATACGAGAAGTCGTCGTGACTGGAGTTCAGACGTGTGCTCTTCCGATCTACTCCCTGATATACCTGCCG | Illumina sequencing reverse primer (hilD P4 promoter and P^tet promote |
| AL09 | CAAGCAGAAGACGGCATACGAGAACTTCGAGTGACTGGAGTTCAGACGTGTGCTCTTCCGATCTACTCCCTGATATACCTGCCG | Illumina sequencing reverse primer (hilD P4 promoter and P^tet promote |
| AL13 | CAAGCAGAAGACGGCATACGAGAAAGGAATGTGACTGGAGTTCAGACGTGTGCTCTTCCGATCTCGCATCCGTATCCACCTGG | Illumina sequencing reverse primer (prgH promoter) |
| AL14 | CAAGCAGAAGACGGCATACGAGAATAGTTGGTGACTGGAGTTCAGACGTGTGCTCTTCCGATCTCGCATCCGTATCCACCTGG | Illumina sequencing reverse primer (prgH promoter) |
| AL15 | CAAGCAGAAGACGGCATACGAGAAGTCGTCGTGACTGGAGTTCAGACGTGTGCTCTTCCGATCTCGCATCCGTATCCACCTGG | Illumina sequencing reverse primer (prgH promoter) |
| AL16 | CAAGCAGAAGACGGCATACGAGAACTTCGAGTGACTGGAGTTCAGACGTGTGCTCTTCCGATCTCGCATCCGTATCCACCTGG | Illumina sequencing reverse primer (prgH promoter) |
| AL17 | CAAGCAGAAGACGGCATACGAGAAGTCGTCGTGACTGGAGTTCAGACGTGTGCTCTTCCGATCTGAGACCAGCCCAGTTTAGCA | Illumina sequencing reverse primer (ompA promoter) |
| AL18 | CAAGCAGAAGACGGCATACGAGAACTTCGAGTGACTGGAGTTCAGACGTGTGCTCTTCCGATCTGAGACCAGCCCAGTTTAGCA | Illumina sequencing reverse primer (ompA promoter) |
| AL21 | TTAGTACTAACGGTCAGGTTGAG | RT primer (hilA) |
| AM24 | CAAGCAGAAGACGGCATACGAGAAAGGAATGTGACTGGAGTTCAGACGTGTGCTCTTCCGATCTCGGGCTCGGGTTTATATGTT | Illumina sequencing reverse primer (hilA promoter) |
| AM25 | CAAGCAGAAGACGGCATACGAGAATAGTTGGTGACTGGAGTTCAGACGTGTGCTCTTCCGATCTCGGGCTCGGGTTTATATGTT | Illumina sequencing reverse primer (hilA promoter) |
| AM26 | CAAGCAGAAGACGGCATACGAGAAGTCGTCGTGACTGGAGTTCAGACGTGTGCTCTTCCGATCTCGGGCTCGGGTTTATATGTT | Illumina sequencing reverse primer (hilA promoter) |
| AM27 | CAAGCAGAAGACGGCATACGAGAACTTCGAGTGACTGGAGTTCAGACGTGTGCTCTTCCGATCTCGGGCTCGGGTTTATATGTT | Illumina sequencing reverse primer (hilA promoter) |
| AL85 | CAAGCAGAAGACGGCATACGAGAAGTCGTCGTGACTGGAGTTCAGACGTGTGCTCTTCCGATCTTGTGTCCTGTGCGGTAATCT | Illumina sequencing reverse primer (hilD P3 promoter) |
| AL86 | CAAGCAGAAGACGGCATACGAGAACTTCGAGTGACTGGAGTTCAGACGTGTGCTCTTCCGATCTTGTGTCCTGTGCGGTAATCT | Illumina sequencing reverse primer (hilD P3 promoter) |
| AM38 | TATAACCTTCCGCCCCGTAC | qPCR forward primer (prgH antisense) |
| AM39 | CAGAGCCCGTCAATTTGTCG | qPCR reverse primer (prgH antisense) |

Red lettering denotes sequences annealing to chromosomal DNA or cDNA in λ red recombineering experiments and in 5'RACE-Seq analysis, respectively. Colored hexameric sequences correspond to indexes used in Illumina sequencing.

### Table S3

**Supplementary Table 3. Relevant alleles constructed by λ red recombineering**

| Allele | Primer pair | template | Notes |
|---|---|---|---|
| hilD-tetR-P^tet-ccdB-cat | ppZ13 - ppZ14 | pNNB5 | AHTc-inducible ccdB on the 3' side of hilD. Step 1 in the construction of markerless hilD-3xFLAG |
| hilD-3xFLAG | ppZ45 - ppZ46 | pSUB11 | |
| hns-3xFLAG_FRT-kan-FRT | ppW78- ppW79 | pSUB11 | |
| ΔhilD::tetRA | AE45 - AE46 | pTn5-tetRA-SH1 | |
| hilAΔK28::FRT-kan-FRT | AF93 - AF26 | pKD13 | FRT-kan-FRT insertion removing a 28266 bp SPI-1 segment from position 338 of hilA to the end of STM2906. Step 1 in the construction of hilA::gfpSF ΔK28_kan |
| ΔhilA338::gfpSF | AG21 - AG39 | pNCM2 | |
| ΔinvB1::gfpSF | AG46 - AG49 | pNCM2 | |
| Δ[hisGDCBHAFIE]::P^Tac-mCherry_FRT-kan-FRT | ppY88 - AA30 | pNCM4 | |
| Δ[sitA-orgA]::tetR-P^tet-ccdB-cat | AE12 - AH30 | pNNB5 | Step 1 in the construction of Δ[sitA-prgH332]::tetR-P^tet |
| Δ[sitA-prgH332]::tetR-P^tet | AH38 - AH39 | Fill-in | Primers anneal to each other |
| Δ[sitA-prgH332]::tetR-P^tet-Term^hisL | AI27 - AI28 | Fill-in | Primers anneal to each other |
| orgB::tetR-P^tet-ccdB-kan | AF19 - AG26 | pNNB7 | Step 1 for moving hilD-3xFLAG in the NusG depletable background |
| hilDΔ309::tetR-P^tet-ccdB-cat | AL78 - AL79 | pNNB5 | Step 1 in the construction of markerless hilD309 |
| hilDΔ309 | AL81 - AL82 | Fill-in | Primers anneal to each other |

### Table S4

**Supplementary Table 4. Primers used in qPCR experiments**

| | | | | |
|---|---|---|---|---|
| Fig. 2a | | | RT primer | qPCR primer pair |
| | gene of interest | hilD | AI41 | AI62 - AI63 |
| | reference gene | ompA | AJ33 | AJ32 - AJ37 |
| Fig. 2b | | | RT primer | qPCR primer pair |
| | gene of interest | prgH | AI48 | AI50 - AI51 |
| | reference gene | ompA | AJ33 | AJ32 - AJ37 |
| Fig. 2c | | | qPCR primer pair | |
| | gene of interest | prgH-hilD IG | AL12 - AI57 | |
| | reference gene | katN | AK19 - AK20 | |
| Fig. 2f | | | RT primer | qPCR-1 primer pair |
| | gene of interest | hilD | AI41 | AI62 - AI63 |
| | reference gene | ompA | AJ33 | AJ32 - AJ37 |
| | | | | qPCR-2 primer pair |
| | gene of interest | prgH antisense | AI41 | AM38 - AM39 |
| | reference gene | ompA | AJ33 | AJ32 - AJ37 |
| Fig. 2g | | | RT primer | qPCR-1 primer pair |
| | gene of interest | prgH | AI48 | AI50 - AI51 |
| | reference gene | ompA | AJ33 | AJ32 - AJ37 |
| | | | | qPCR-2 primer pair |
| | gene of interest | hilD antisense | AI48 | AI62 - AI63 |
| | reference gene | ompA | AJ33 | AJ32 - AJ37 |
| Fig. 3b | | | RT primer | qPCR primer pair |
| | gene of interest | hilD | AI41 | AI62 - AI63 |
| | reference gene | ompA | AJ33 | AJ32 - AJ37 |
| Fig. 3c | | | RT primer | qPCR primer pair |
| | gene of interest | prgH | AI48 | AI50 - AI51 |
| | reference gene | ompA | AJ33 | AJ32 - AJ37 |
| Fig. 3f | | | qPCR primer pair | |
| | gene of interest | prgH-hilD IG | AL12 - AI57 | |
| | reference gene | katN | AK19 - AK20 | |
